# Unveiling the guardians: IL-26-expressing MAIT cells protect epithelial barrier function and are dysregulated in Crohn’s disease

**DOI:** 10.1101/2024.06.02.597015

**Authors:** Veronika Bosáková, Bo-Jun Ke, Marcela Hortová Kohoutková, Ioanna Papatheodorou, Filip Kafka, Marco De Zuani, Sneha Santhosh, Francesca Biscu, Saeed Abdurahiman, Ine De Greef, Sare Verstockt, Bram Verstockt, Séverine Vermeire, Rafael J Argüello, Gianluca Matteoli, Jan Frič

**Affiliations:** International Clinical Research Center, St. Anne’s University Hospital Brno, Czech Republic; Department of Biology, Faculty of Medicine, Masaryk University, Brno, Czech Republic; Translational Research Center for Gastrointestinal Disorders (TARGID), Department of Chronic Diseases and Metabolism, KU Leuven, 3000 Leuven, Belgium; International Clinical Research Center, Faculty of Medicine, Masaryk University, Brno, Czech Republic; Centre for Inflammation Research, University of Edinburgh, Edinburgh, UK; Department of Gastroenterology and Hepatology, University Hospitals Leuven, Leuven, Belgium; Leuven Institute for Single-cell Omics (LISCO), KU Leuven, Leuven, Belgium; Aix Marseille Univ, CNRS, INSERM, CIML, Centre d’Immunologie de Marseille-Luminy, Marseille, France; Institute of Hematology and Blood Transfusion, Prague, Czech Republic

**Keywords:** IBD, intestinal inflammation, mucosal-associated invariant T cells, IL-26, iPSC-derived organoids

## Abstract

**BACKGROUND & AIMS:** Inflammatory bowel disease (IBD) is characterized by a dysregulated immune response against the host’s microbiome. Mucosal-associated invariant T (MAIT) cells recognize microbiota-derived riboflavin metabolites and play a crucial role in mucosal homeostasis. However, their specific role in IBD remains enigmatic. MAIT cells express IL-26, a novel IL-10 family cytokine with a controversial role in IBD. We investigated the functions of MAIT cells and IL-26 in IBD using a unique combination of state-of-the-art 3D human intestinal tissue models and clinical samples.

**METHODS:** We analyzed MAIT cells from the peripheral blood and intestinal tissue of Crohn’s disease (CD) patients, using immunofluorescence staining and flow cytometry to describe the phenotype and IL-26 expression of MAIT cells. We used 3D iPSC-derived intestinal organoids as a complex *in vitro* model of human tissue and RNA sequencing and functional assays such as wound healing assay to study the role of IL-26 in mucosal homeostasis and inflammation.

**RESULTS:** We observed a reduction of MAIT cells in the peripheral blood of CD patients compared to healthy donors (1.5 ± 0.4%; 4.1 ± 1.1%; p < .0065) and a significant decrease of MAIT cells in inflamed compared to non-inflamed ileum of CD patients (0.1 ± 0.03%; 0.17 ± 0.05%; p < .042). MAIT cells were found pathologically activated in inflamed tissue, exhibiting differences in CD8 and CD4 expression and dysregulation of IL-26 expression. Furthermore, we demonstrated a protective role of IL-26 in mucosal homeostasis and inflammation in the iPSC-derived organoid model.

**CONCLUSION:** Our results show a crucial role for IL-26 and MAIT cells in the homeostasis of intestinal tissue and in the pathogenesis of IBD. These cells may therefore represent new therapeutic targets for CD patients.

## INTRODUCTION

Inflammatory bowel disease (IBD), including Crohn’s disease (CD) and ulcerative colitis (UC), is a progressive disorder affecting the gastrointestinal tracts of millions of people worldwide. Despite extensive research, the exact cause of IBD remains unknown, while incidence is still increasing^1^. In general, IBD is considered the result of a dysregulated immune response targeting the gut microbiome. Unconventional T cells, including mucosal-associated invariant T (MAIT) cells, bridge innate and adaptive immunity to orchestrate a proper response to microorganisms – therefore, understanding their role in mucosal immunity is critical for unraveling the pathogenesis of IBD.

MAIT cells represent 1 - 10% of all T cells in human peripheral blood^2, 3^ and up to 10% of T cells in mucosal tissues including the intestine^3, 4^. They express a highly evolutionarily conserved invariant ⍺-chain of the T-cell receptor (TCR Vα7.2)^5^ recognizing MHC class I-related molecule 1 (MR1), which is ubiquitously expressed in various cell types and tissues^6^. MAIT cells are activated by metabolites of the microbial riboflavin synthesis pathway in a TCR-dependent manner^7, 8^. IBD is characterized by profound changes in microbiome composition including a loss of commensals, and a significant decrease in microbial diversity and richness^9^. Intriguingly, high diversity in the microbiome is associated with low activation potential of MAIT cells and higher demand for riboflavin, and *vice versa*^10^. Several studies have implicated MAIT cells in the pathophysiology of IBD^11–16^, but their exact mechanism of action remains unclear.

MAIT cells respond rapidly after activation, expressing a mixed pattern of Th1 and Th17 cytokines including IL-26^17–19^, a recently discovered cytokine belonging to the IL-10 family. Interestingly, IL-26 is not expressed in rodents – therefore, its effects on inflammation have not been fully investigated *in vivo*^20, 21^, with some exceptions where humanized mice were used to explore the role of IL-26 in the progression of induced colitis^22^. This study showed a protective role for IL-26, and IL-26-expressing mice had milder symptoms of acute colitis and decreased expression of proinflammatory cytokines in intestinal tissue^22^. Single-nucleotide polymorphisms in the gene encoding IL-26 have also been associated with UC^23^, further suggesting a role in the pathology of IBD.

While IBD has been extensively studied in mice, these models fall short in replicating the role of human MAIT cells in intestinal inflammation^20, 21^. This discrepancy is amplified by the absence of IL-26 expression in mice, significantly impacting the study of this cytokine’s role in IBD. To overcome these problems, organoids have been employed as a 3D *in vitro* model of the human gut that maintains *in vivo*-like tissue complexity. We have previously shown that intestinal organoids (IOs) form an immunocompetent microenvironment^25^ and respond to stimulation by various pathogen pattern recognition receptor ligands with a significant upregulation of proinflammatory chemokines and cytokines^25^. They also form complex 3D structures that are highly organized and physiologically mimic human intestinal tissue. Therefore, they represent a unique human *in vitro* model to study the roles of MAIT cells and IL-26 in mucosal tissue. Recently, human induced pluripotent stem cell (hiPSC)-derived organoids were used to model a UC-like milieu *in vitro*, illustrating the potential of hiPSC-derived organoids as models of mucosal inflammation and as a drug testing tool offering opportunities for personalized medicine^26^.

Here, we show that MAIT cells are less frequent in the peripheral blood and ileum of CD patients and acquire a distinct phenotype compared to healthy donors. We used 3D human iPSC-derived IOs to model healthy and proinflammatory environments, allowing us to demonstrate a protective role of IL-26 under both conditions and therefore unravel its therapeutic potential in IBD. The results show that MAIT cells and IL-26 play important roles in the protection of intestinal homeostasis and may suggest them as new therapeutic targets for IBD.

## Materials and Methods

### Human specimens

Resected terminal ileum was collected immediately following ileocaecal resection from patients with Crohn’s disease (CD) under the supervision of the specialized IBD pathologist. All blood samples were collected in the EDTA tube from CD patients one day before surgery. The protocol was approved by the Institutional Review Board (IRB) of the University Hospitals Leuven, Belgium (B322201213950/S53684, CCARE, S-53684). All recruitment was performed after ethical approval and oversight from the IRB and informed consent was obtained from all participants before surgery. Clinical information and metadata for the patients included in this study are provided in **Table S1**.

### Flow cytometry analysis

Single-cell suspensions obtained from intestinal tissue and peripheral blood mononuclear cells (PBMCs) were used for flow cytometry. Firstly, the cells were stained with Fixable Viability Dye eFluor™780 (1:1000, eBioscience). After washing, the cells were stained with fluorochrome-conjugated anti-human CD45, CD3, CD4, CD8, CD14, CD19, CD69, CD161, and TCR Vα7.2 antibodies. After staining, the cells were fixed and permeabilized using the eBioscience IC staining kit (eBioscience). The cells were stained with IL-26 antibody (all antibody details are shown in **Table S2**). The MAIT cells were identified as the Live/death^-^ CD45^+^Lin(CD14, CD19)^-^CD3^+^CD161^+^TCR Vα7.2^+^ cell population (gating strategy **Figure S1A, B**). The samples were analyzed on an ID7000 spectral flow cytometer (Sony). Data were analyzed using FlowJo v10 software (BD Life Sciences).

### Intestinal organoids (IOs) differentiation

IOs were differentiated from human induced pluripotent stem cells (hiPSCs) using an adapted protocol^25, 27^. In brief, hiPSCs were differentiated to definitive endoderm (DE) using Activin A (100 ng/mL, R&D) in RPMI-1640 (Gibco) with increasing concentrations of HyClone-defined fetal bovine serum (HC FBS) (GE Healthcare Bio-Sciences) (first day 0%, second day 0.2%, and third and fourth days 2%). Next, the DE was differentiated to mid- and hindgut using RPMI-1640 supplemented with 15 mM HEPES, 2% HC FBS, FGF-4 (500 ng/mL, R&D), and WNT-3a (500 ng/mL, R&D) until 3D spheroids were observed. Spheroids were collected and seeded in Cultrex Membrane Extract, Type 2 (R&D). The spheroids were fed with IOs medium (Advanced DMEM F12 (Gibco) supplemented with 15 mM HEPES, B27 supplement (Thermo Fisher Scientific), GlutaMAX supplement (Thermo Fisher Scientific), P/S (500 U/mL), R-Spondin 1 (500 ng/mL, R&D), Noggin (100 ng/mL, R&D), and EGF (100 ng/mL, R&D)). The medium was changed twice a week. IOs that were at least 50 days old were used for the experiments.

### Statistical analyses

Statistical analyses were performed with GraphPad Prism 9. Data were tested for normality prior to the use of parametric or nonparametric tests. The normality was tested using the Shapiro-Wilk test. Depending on the dataset, paired/unpaired *t*-test, Wilcoxon test, Mann-Whitney test, or one-way ANOVA with Tukey’s multiple comparison test was used. Data are shown as mean ± SEM unless otherwise stated. In violin plots, dashed lines represent median and dotted lines represent quartiles.

**Immunofluorescence staining of tissue samples, isolation of single-cell suspension from the ileum, peripheral blood mononuclear cells isolation, MAIT cells isolation, *in vitro* activation of MAIT cells, intracellular staining for *in vitro* expression of IL-26, Annexin V staining, Single-cell metabolism analysis (SCENITH^TM^), RNA extraction and gene expression analysis, human induced pluripotent stem cells maintenance, dissociation of IOs, Immunofluorescence staining for IOs, stimulation of the IOs, bead array cytokines detection, RNA sequencing, western blot, wound healing assay, apoptosis assay for IOs, co-cultivation of MAIT cells with IOs.**

See **Supplementary Materials and Methods**, **Supplementary table 1**, **Supplementary table 2** and **Supplementary table 3** for details.

## RESULTS

### MAIT cells are diminished in peripheral blood and ileum of CD patients

To detect disease-associated changes in frequency and phenotype, we isolated MAIT cells from inflamed (INF) and non-inflamed (NON-INF) tissues obtained from biopsies of CD patients (**Figure 1A**) (**Table S1**). We compared INF and NON-INF tissue from the same patient’s ileocecal resections, using immunofluorescence staining to identify and localize TCR Vα7.2^+^CD161^+^ MAIT cells within the intestinal tissue. A clear decrease of MAIT cells was observed in the INF compared to NON-INF tissue (**Figure 1B-D**). In both conditions, MAIT cells were localized in close proximity to intestinal crypts. We next quantified the relative frequency of MAIT cells using flow cytometry (**Figure S1A, B**), showing a significant decrease of MAIT cells among live cells in INF (0.1 ± 0.03%) compared to NON-INF (0.17 ± 0.05%) tissue (**Figure 1E**). Furthermore, MAIT cells collected from the peripheral blood of the CD patients prior to surgery (**Figure 1F**) demonstrated a significant decrease of MAIT cells among the CD3^+^ cells (**Figure 1G**) and CD45^+^ cells (**Figure 1H**) in CD patients (1.5 ± 0.4%; 0.6 ± 0.2%) compared to healthy donors (HD) (4.1 ± 1.1%; 2.4 ± 0.6%). Importantly, the percentages of live cells (**Figure S1C**) and CD3^+^ cells (**Figure S1D**) were comparable in INF compared to NON-INF tissue and in the peripheral blood of CD patients compared to HD. **Overall, our results show a significant decrease of MAIT cells in both the peripheral blood and inflamed tissue of CD patients.**

**Figure 1.**
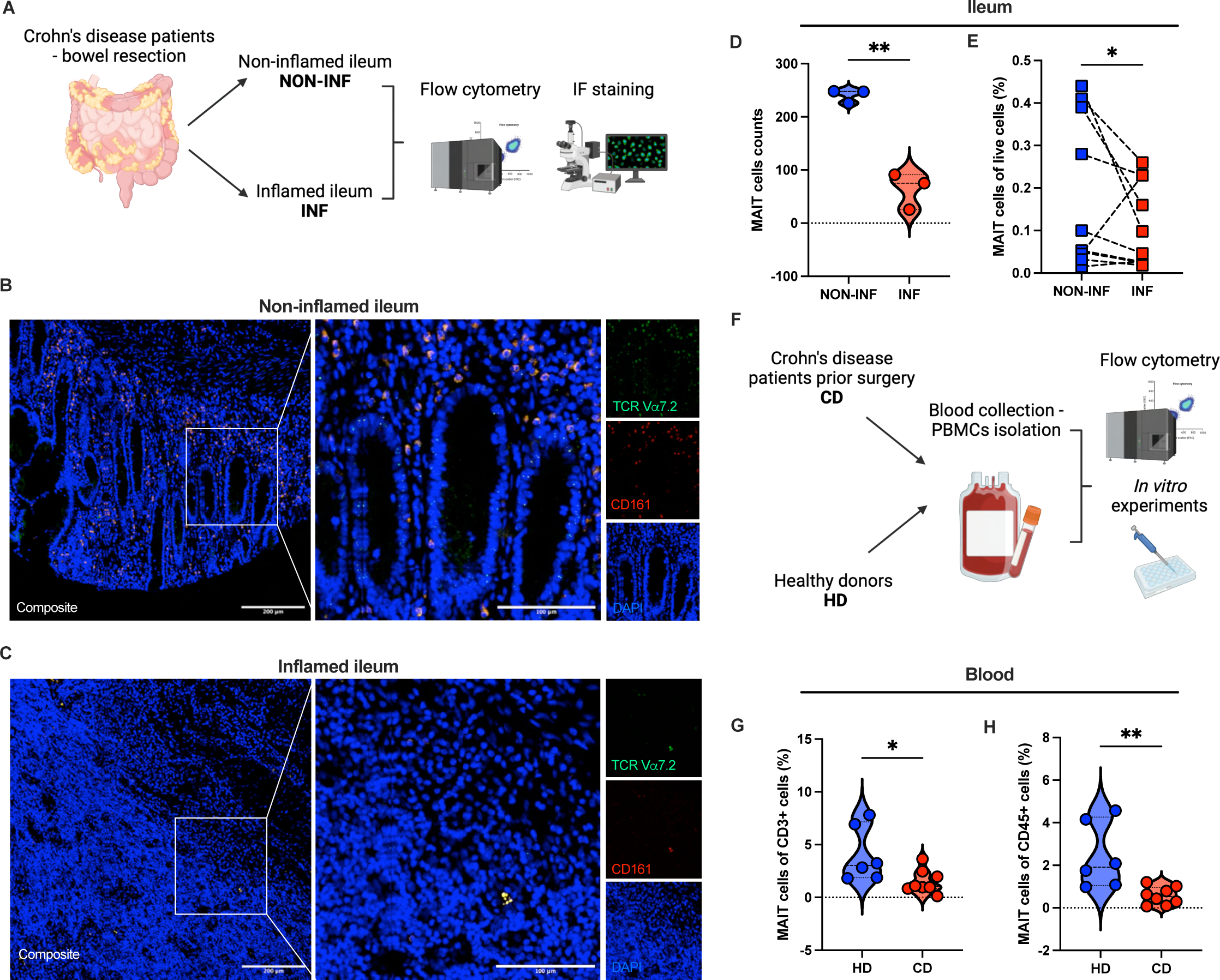
The abundance of MAIT cells is decreased in CD patients. (A) Workflow for tissue sample processing. (B, C) Immunofluorescence staining (25x magnification) of NON-INF (B) and INF tissue (C). (D) Quantification of MAIT cells in immunofluorescence staining. (E) Percentages of MAIT cells among live cells from tissue samples obtained using flow cytometry. Dashed lines connect paired data points from the same patient. (F) Schematic of peripheral blood samples processing. (G, H) Percentages of MAIT cells among CD3^+^ (G) and CD45^+^ cells (H) in peripheral blood. \**p* < .05, \*\**p* < .01.

### MAIT cells display an activated phenotype in CD patients

Next, we analyzed phenotypic changes in MAIT cells under homeostasis and inflammatory conditions in CD patients. We used flow cytometry (**Figure 2A**) for patient sample analysis and *in vitro* and *ex vivo* experiments (**Figure 2B**) to further describe the MAIT cell population. First, we assessed the activation status of MAIT cells by comparing the expression of the activation marker CD69 in INF and NON-INF tissue and peripheral blood from CD patients and HD. We observed a significantly higher percentage of CD69^+^ MAIT cells in INF (16.5 ± 4.6%) compared to NON-INF tissue (4.7 ± 1.5%) (**Figure 2C**). CD69^+^ MAIT cells were similarly increased in peripheral blood from CD patients (23.7 ± 6.7%) compared to HD (6.0 ± 2.4%) (**Figure 2D**). In order to better understand these physiological changes, we sought to replicate the observed MAIT cell activation *in vitro.* FACS-sorted MAIT cells from peripheral blood mononuclear cells (PBMCs) of HD were stimulated using anti-CD3 and -CD28-coated beads to mimic TCR-dependent (TCRd) activation. In addition to TCRd stimulation, MAIT cells can be activated through the fully TCR-independent (TCRi) pathway by cytokines. To mimic TCRi activation, we used a combination of MAIT cell-activating cytokines including IL-12, IL-15, IL-18, and TL1A^8, 28, 29^ (**Figure 2B**). We first assessed CD69 expression upon activation, finding a significant increase of CD69^+^ MAIT cells after all types of stimulation (**Figure S2A**). Next, we used Annexin V staining to identify apoptotic cells. The subsequent analysis revealed a significantly higher percentage of Annexin V^+^ MAIT cells after TCRi and TCRd+TCRi but not after TCRd activation (**Figure 2E left**).

**Figure 2.**
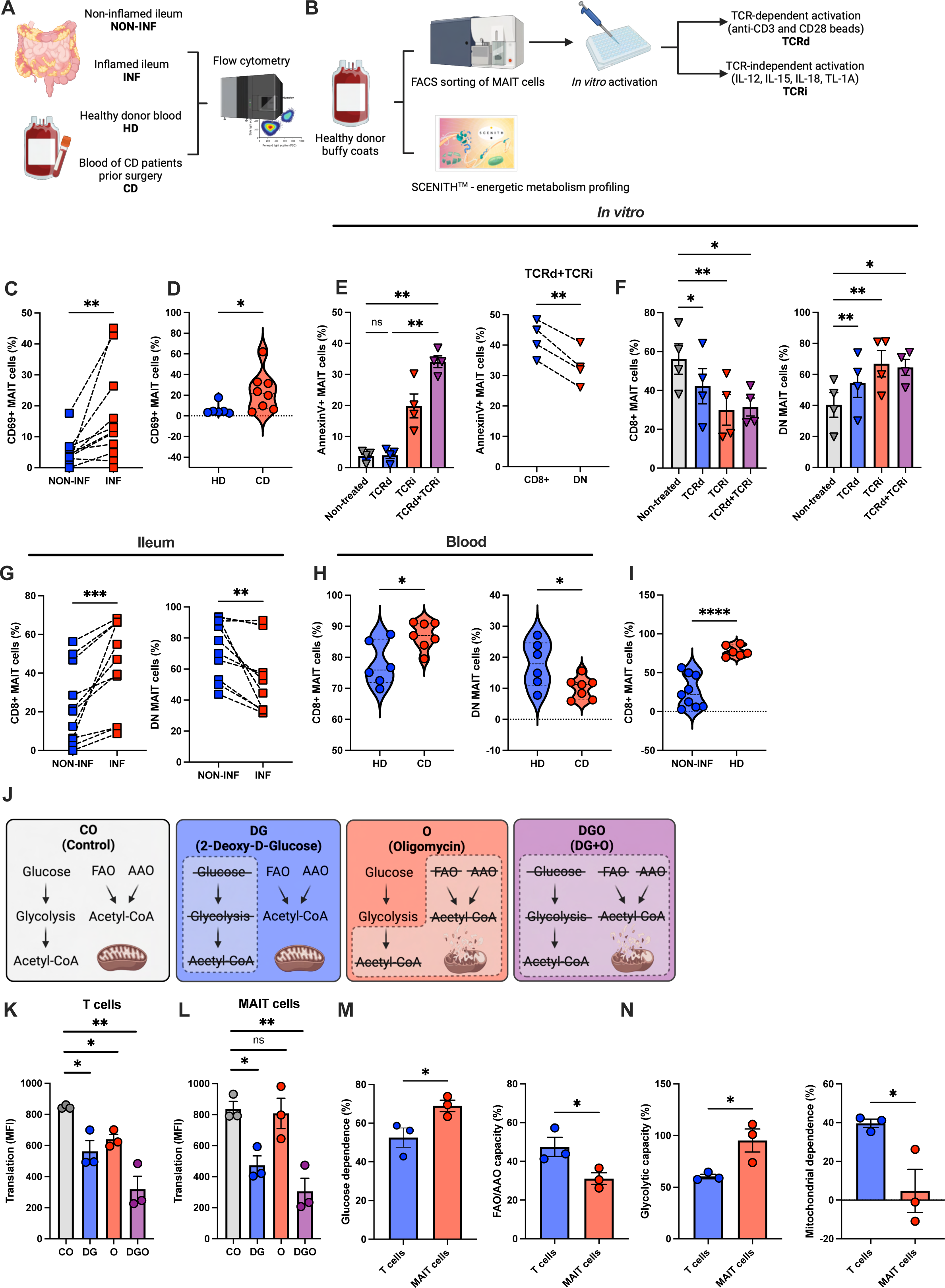
MAIT cells exhibit a distinct phenotype in CD patients. (A, B) Schematic for patient samples processing (A) and *in vitro* experiments performed using FACS-sorted MAIT cells (B). (C, D) Percentage of CD69^+^ MAIT cells in the ileum (C) and peripheral blood (D). (E) Percentage of Annexin V^+^ MAIT cells after 72 hours of *in vitro* stimulation. (F-H) The proportion of CD8^+^ and DN MAIT cells after 72 hours of *in vitro* stimulation (F), in INF and NON-INF tissue (G), and in peripheral blood (H). (I) Violin plot showing the percentage of CD8^+^ MAIT cells in NON-INF tissue and peripheral blood from HD. (J) Scheme of SCENITH^TM^. (K, L) Energetic metabolism of conventional T cells (K) and MAIT cells (L) from peripheral blood of HD at steady state. (M) Relative glucose dependence (left) and FAO/AAO capacity (right) of MAIT cells and conventional T cells from peripheral blood of HD at steady state. (N) Relative glycolytic capacity (left) and mitochondrial dependence (right) in MAIT cells and conventional T cells from peripheral blood of HD at steady state. For plots (C, E right, G) dashed lines indicate paired data points. \**p* < .05, \*\**p* < .01, ns, not significant.

MAIT cells can be divided into distinct subsets based on their expression of CD4 and CD8. Therefore, further analysis was focused on the characterization of CD4^-^CD8^+^ (CD8^+^) and CD4^-^ CD8^-^ (DN) MAIT cells at steady state and after *in vitro* activation. We observed no significant differences in CD69 expression by CD8^+^ and DN MAIT cells at steady state and after TCRd and TCRd+TCRi activation (**Figure S2B**); however, CD69 expression was significantly higher in CD8^+^ vs. DN MAIT cells after TCRi stimulation (**Figure S2B**). CD8^+^ MAIT cells also displayed more proapoptotic features compared to DN MAIT cells after TCRi and TCRd+TCRi activation (**Figure 2E right, S2C right**), while no differences were observed in Annexin V^+^ CD8^+^ and DN MAIT cells at steady state or after TCRd activation (**Figure S2C left and middle**).

Notably, our analysis showed a significant decrease in CD8^+^ (**Figure 2F left**) and increase in DN (**Figure 2F right**) MAIT cells after all types of *in vitro* activation. We also analyzed these percentages in the ileum of CD patients. These results showed the opposite trend: an increase of CD8^+^ (**Figure 2G left**) and a decrease of DN (**Figure 2G right**) MAIT cells in INF compared to NON-INF tissue. Decreased CD8^+^ (**Figure 2H left**) and increased DN (**Figure 2H right**) MAIT cells were also observed in the peripheral blood of CD patients compared to HD. Interestingly, the percentage of CD8^+^ MAIT cells was significantly lower in NON-INF tissue of CD patients compared to peripheral blood of HD (**Figure 2I**), suggesting different functionality of MAIT cells in context of tissue and peripheral blood. **Altogether, MAIT cells display a more activated phenotype in the peripheral blood and inflamed tissue of CD patients, and CD4 and CD8 expression by MAIT cells differs between CD patients and HD.**

### MAIT cells form a metabolically separate population from conventional T cells

Energy metabolism is critical for immune functionality; therefore, we sought to address the metabolic requirements of MAIT cells in order to understand their possible dysregulation in CD patients. We used SCENITH^TM^, a method allowing single-cell, flow cytometric profiling of energy metabolism by monitoring changes in protein translation upon metabolic pathway inhibition^30^ (**Figure 2J**), to assess the energy metabolism profiles of MAIT cells and conventional T cells from peripheral blood of HD. In conventional T cells, we observed a significant decrease of translation when glycolysis and fatty and amino acid oxidation (FAO/AAO) were inhibited using 2-deoxy-D-glucose (DG) and oligomycin (O), respectively (**Figure 2K**). However, translation in MAIT cells was affected only when glycolysis was inhibited (**Figure 2L**). Further analysis showed that MAIT cells exhibited significantly higher glucose dependence (**Figure 2M left**), correspondingly lower FAO/AAO capacity (**Figure 2M right**), and significantly higher glycolytic capacity and lower mitochondrial dependence (**Figure 2N**) compared to conventional T cells. After TCRd+TCRi activation, the metabolic profile of MAIT cells changed, and the differences between them and conventional T cells disappeared (**Figure S2D**). **Overall, MAIT cells are a metabolically distinct T cell subpopulation with high glucose dependence and glycolytic capacity.**

### IL-26 expression by MAIT cells is limited to ileum and is dysregulated in CD patients

As MAIT cells are known to express IL-26^17–19^, we aimed to confirm the expression of IL-26 in human ileum. Immunofluorescence staining of NON-INF tissue from CD patients showed expression of IL-26 mainly by CD161^+^ cells in close proximity to intestinal crypts (**Figure 3A**). We performed flow cytometry to address the expression of IL-26 by MAIT cells, finding that IL-26^+^ MAIT cells were present in NON-INF tissue from CD patients (**Figure 3B, C**). Interestingly, a comparison of INF and NON-INF tissue revealed a significant decrease of IL-26^+^ MAIT cells (13.3 ± 3.96%; 21.4 ± 5.2%) under inflammatory conditions (**Figure 3B, D**). Next, we focused on IL-26 production by MAIT cells in peripheral blood of CD patients and HD. The flow cytometry analysis revealed almost no IL-26^+^ MAIT cells in peripheral blood of HD (0.08 ± 0.02%) (**Figure 3E, F**) and significantly higher levels of IL-26^+^ MAIT cells in CD patients (1.2 ± 0.4%) (**Figure 3E, G**).

**Figure 3.**
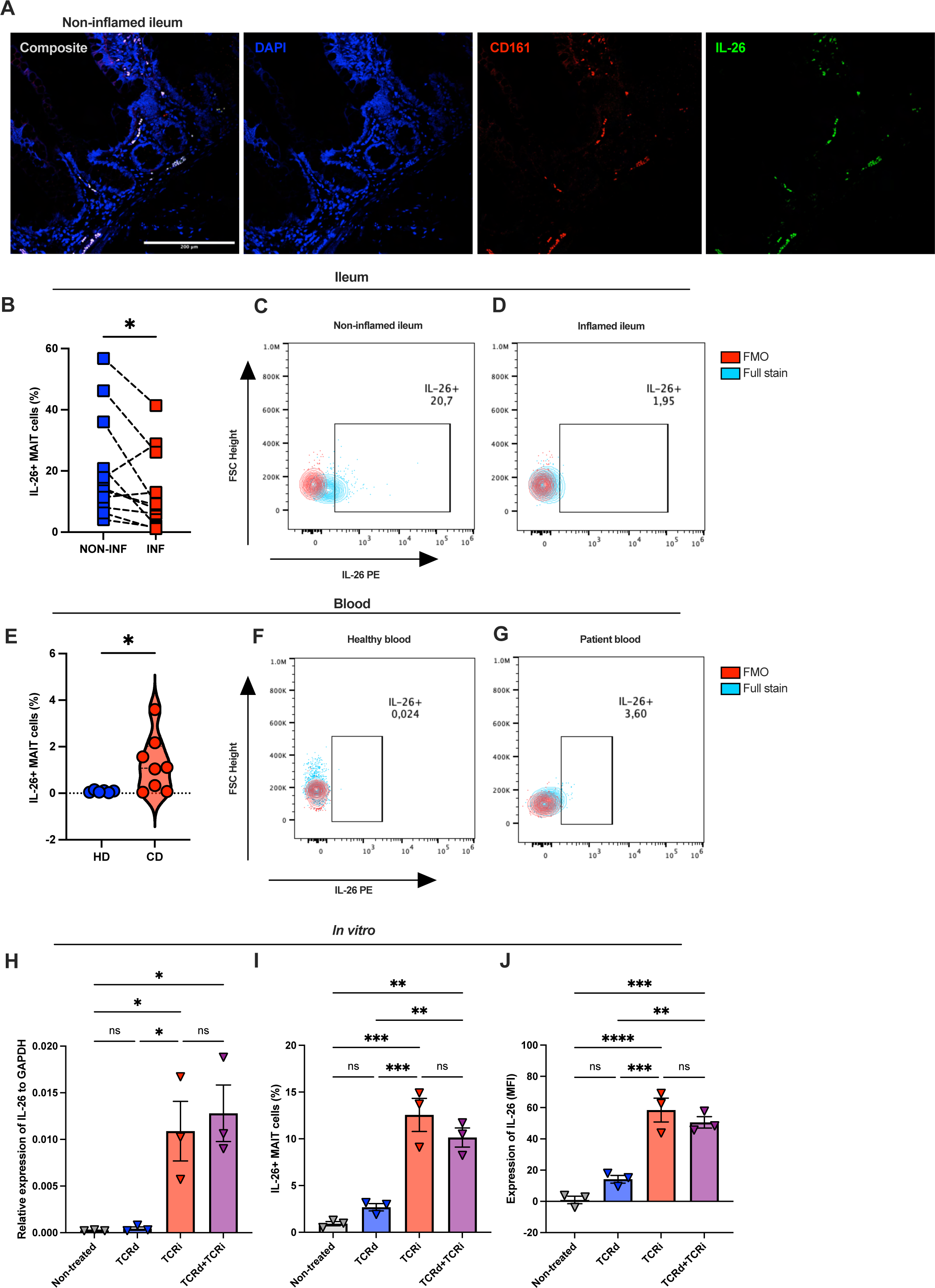
IL-26 expression by MAIT cells is restricted to intestinal tissue. (A) Immunofluorescence image (25x magnification) of NON-INF intestinal tissue. (B) Proportion of IL-26^+^ MAIT cells in NON-INF and INF tissue. Dashed lines connect paired datapoints from the same patient. (C, D) Contour plots showing IL-26 expression in MAIT cells in NON-INF (C) and INF tissue (D). (E) Percentage of IL-26^+^ MAIT cells in peripheral blood. (F, G) Contour plots showing IL-26 expression in MAIT cells in peripheral blood from HD (F) and CD patients (G). (H) IL-26 RNA expression in MAIT cells after *in vitro* stimulation. (I, J) Plots showing the percentage of IL-26^+^ MAIT cells (I) and IL-26 expression in MAIT cells (J) after the indicated *in vitro* stimulation regimens. \**p* < .05, \*\**p* < .01, \*\*\**p* < .005, \*\*\*\**p* < .001, ns, not significant.

To further describe the pathways involved in IL-26 production, we activated FACS-sorted MAIT cells from HD with TCRi, TCRd, or TCRd+TCRi. Expression of IL-26 RNA was significantly induced only upon TCRi/TCRd+TCRi activation (**Figure 3H**), and further flow analysis demonstrated a similar trend on the protein level (**Figure 3I, J**). No significant differences were observed in IL-26 expression between CD8^+^ and DN MAIT cells (**Figure S3A**). **In conclusion, our results show that TCRi activation of MAIT cells is critical for their production of IL-26. Moreover, under non-inflammatory conditions, IL-26 expression is restricted to intestinal tissue. In CD patients, this pattern is dysregulated: IL-26 expression is significantly upregulated in peripheral blood and downregulated in the ileum.**

### IL-26 drives differential expression of genes involved in tight junction assembly and maintenance of intestinal tissue homeostasis

After demonstrating that IL-26 is produced by intestinal tissue MAIT cells, we focused on describing its influence on the expression of proinflammatory mediators and molecules involved in tissue homeostasis. Human induced pluripotent stem cell (hiPSC)-derived intestinal organoids (IOs), were used to model healthy human intestinal tissue, as they allow to study effect on both epithelial and mesenchymal cells in complex 3D model. These IOs express both domains of the IL-26 receptor, IL-10Rβ and IL-20Rɑ, in epithelial and mesenchymal cells (**Figure 4A, S4A-C**). Stimulation of IOs with human recombinant (hr)IL-26 induced a significant increase of STAT3 phosphorylation (**Figure S4D, E**) but did not induce any significant changes in the RNA (**Figure S4F**) or protein (**Figure S4G**) expression levels of major proinflammatory cytokines and chemokines.

**Figure 4.**
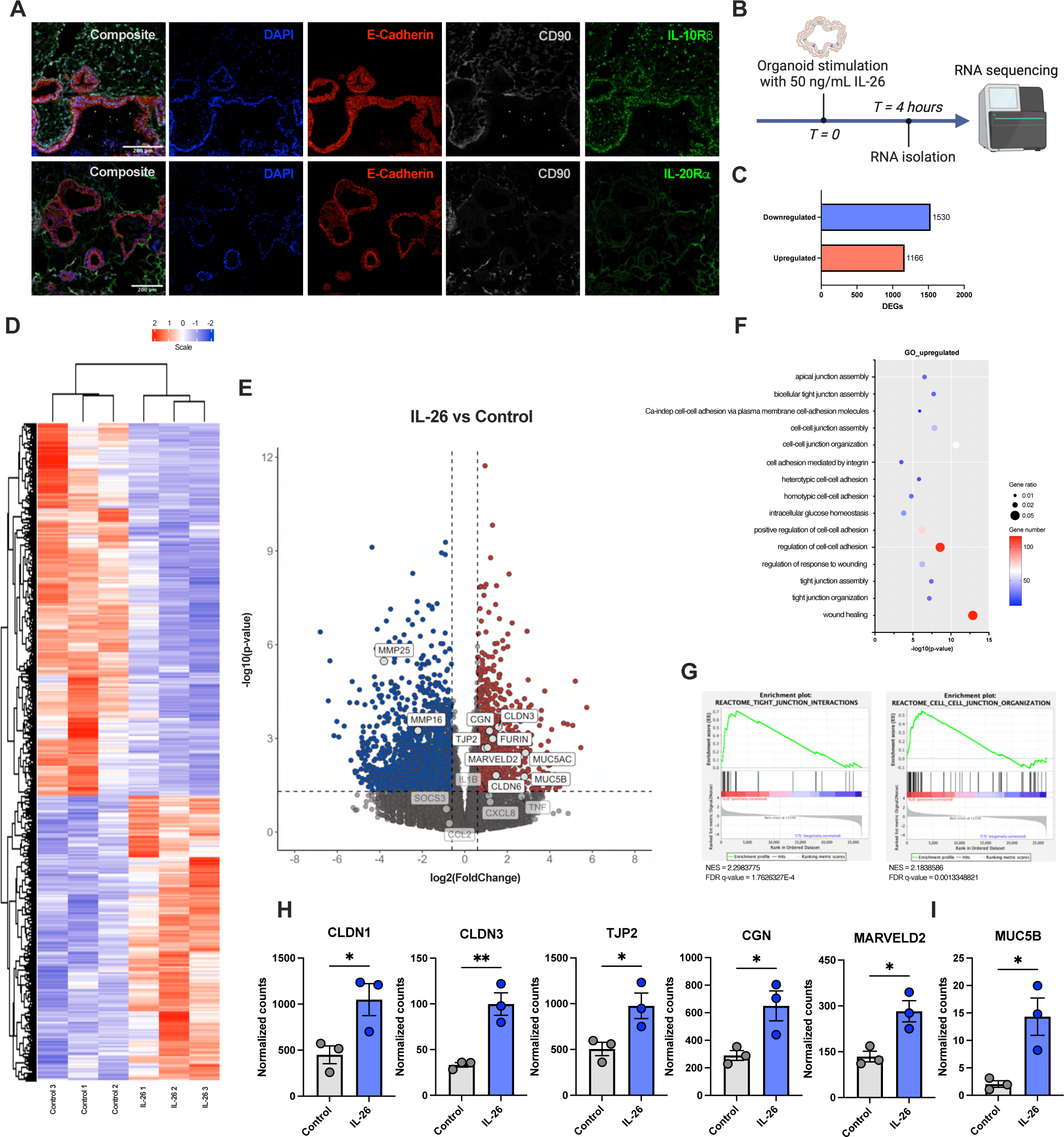
IL-26 drives changes in the expression of genes involved in maintaining intestinal tissue homeostasis. (A) Immunofluorescence images of IOs showing IL-10Rβ and IL-20Rα expression (10x magnification). (B) Schematic of *in vitro* IL-26 stimulation. (C) DEGs in IL-26-treated IOs versus control. (D) Heatmap showing relative expression of genes affected by IL-26 stimulation and clustering of samples. (E) Volcano plot depicting DEGs in IL-26-stimulated compared to untreated IOs. Dots represent significantly upregulated (red) or downregulated (blue) DEGs with *p* < .05. Vertical dashed lines show the |log2(FoldChange)| = 0.6; horizontal dashed line shows *p* = .05. (F) Bubble chart showing the most significantly enriched pathways obtained using gene ontology. (G) Enrichment plots of GSEA. (H, I) Plots showing gene expression using normalized counts. \**p* < .05, \*\**p* < .01; NES, normalized enrichment score; FDR, false discovery rate.

Aiming to describe the overall effects of IL-26 using a 3D *in vitro* model of healthy intestinal tissue, we performed RNA sequencing (RNA-seq) on IL-26-stimulated hiPSC-derived IOs (**Figure 4B**). Differentially expressed genes (DEGs) analysis revealed 2,696 significantly altered genes (**Figure 4C**), and untreated and IL-26-treated samples clustering indicated consistent and specific responses of IOs to IL-26 (**Figure 4D**). We observed significant upregulation of genes such as FURIN, CLDN16, and MUC5AC and downregulation of MMP25 and MMP16 in IL-26-treated samples (**Figure 4E**). Consistent with our previous results (**Figure S4F**), IL-26 stimulation did not affect the expression of major proinflammatory cytokines and chemokines (**Figure 4E**). Functional enrichment analysis of gene ontology using clusterProfiler identified gene pathways affected by IL-26 including *cell-cell junction organization*, *tight junction assembly*, and *apical junction assembly* (**Figure 4F**). Similarly, Gene Set Enrichment Analysis (GSEA) showed enrichment in the *tight junction interactions* and *cell-cell junction organization* pathways (**Figure 4G**). As an intact epithelial barrier is crucial for intestinal tissue homeostasis, we focused our analysis on genes involved in the maintenance of this barrier. We found that IL-26 increased the expression of genes involved in tight junction assembly (including CLDN1, CLDN3, TJP2, CGN, and MARVELD2; **Figure 4H**) and in mucus layer formation (MUC5B; **Figure 4I**). **Overall, our results showed that IL-26 is involved in the processes protecting epithelial barrier function and intestinal tissue homeostasis.**

### IL-26 decreases expression of proinflammatory cytokines and chemokines induced by TNFα in organoid model

Next, to study the CD gut environment and the role of IL-26 under inflammatory conditions, we stimulated IOs with TNFα in the presence or absence of IL-26, and gene expression changes were analyzed using RNA-seq. This analysis identified 1,508 unique DEGs in TNFα+IL-26-treated samples vs. TNFα- or IL-26-treated samples when compared to control samples (**Figure 5A**). 502 and 756 genes were upregulated (**Figure 5B**) and downregulated (**Figure 5C**), respectively. In order to depict the role of IL-26 in proinflammatory condition, we analyzed DEGs in TNFα+IL-26-treated samples compared to TNFα (**Figure 5D**). Further analysis of RNA-seq data revealed that IL-26 stimulation decreased the expression of the IL11, IL15, CCL7, CCL11, CXCL9, and CXCL10 genes (**Figure 5D, S5A, B**).

**Figure 5.**
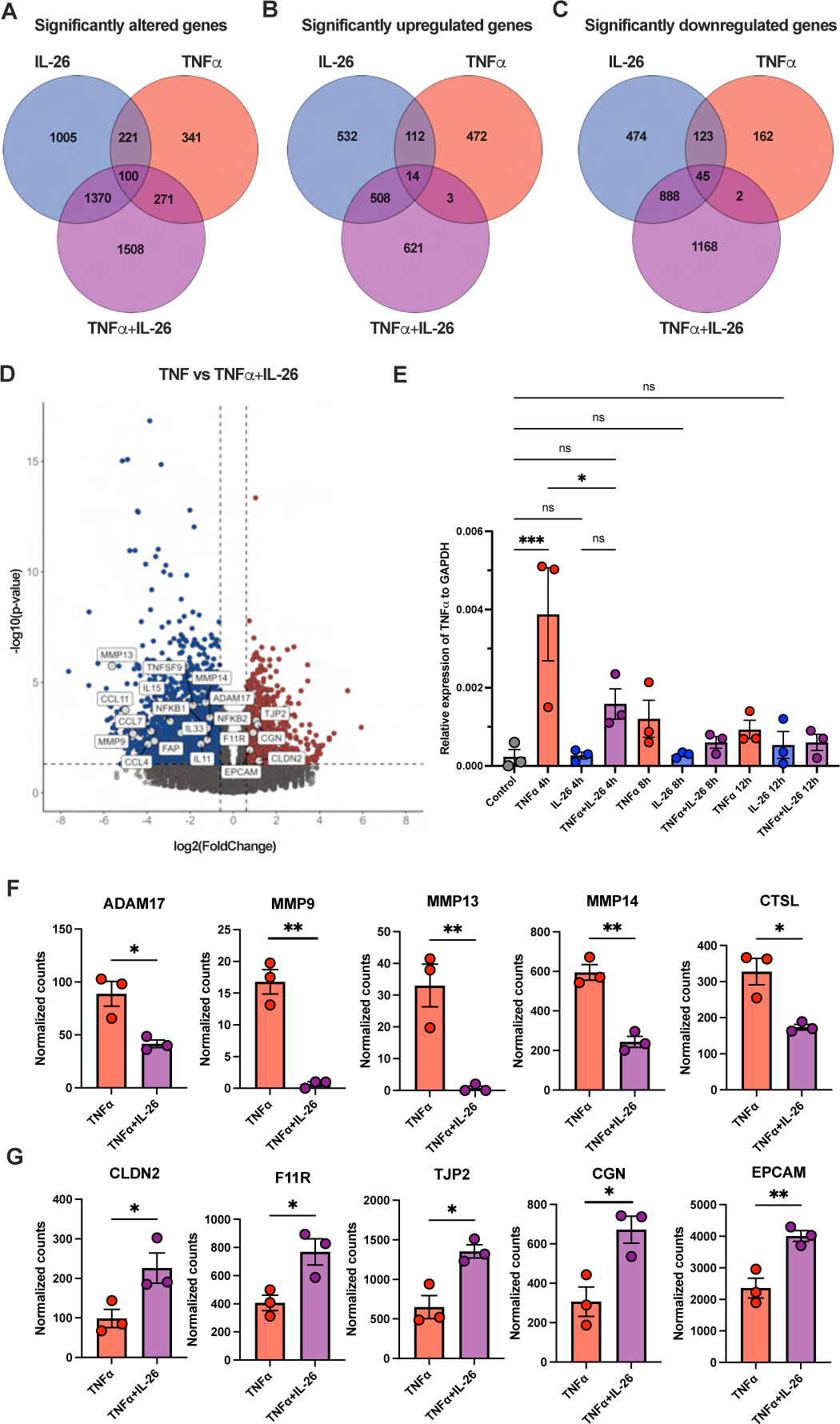
IL-26 has an immunoregulatory role in proinflammatory conditions. (A-C) Venn diagrams depicting total (A), upregulated (B), and downregulated (C) DEGs in IL-26-, TNFα-, and TNFα+IL-26-treated samples compared to control. (D) Volcano plot showing DEGs in TNFα+IL-26-treated samples compared to TNFα-treated samples. Dots represent significantly upregulated (red) or downregulated (blue) DEGs with *p* < .05. Vertical dashed lines show the |log2(FoldChange)| = 0.6; horizontal dashed line shows *p* = .05. (E) TNF gene expression induced by different treatments of IOs at different time points. (F, G) Plots showing gene expression using normalized counts. \**p* < .05, \*\**p* < .01, \*\*\**p* < .005, ns, not significant.

Interestingly, TNFα induced upregulation of TNF gene expression at the RNA level, and this upregulation was decreased when IL-26 was present (**Figure 5E**). As TNFα production can be regulated on the protein level by enzymatic cleavage of its precursor, we also focused on the expression of genes encoding enzymes capable of cleaving pro-TNF^31–34^. We found that IL-26 induced significant downregulation of the expression of genes such as ADAM17, MMP9, MMP13, MMP14, and CTSL (**Figure 5F**). Additionally, having previously observed that IL-26 affects the expression of genes involved in tight junctions and epithelial barrier function in healthy intestinal tissue, we examined these pathways in our model of inflamed gut as well. The expression of genes such as CLDN2, F11R, TJP2, CGN, and EPCAM was significantly increased in TNFα+IL-26-treated samples compared to TNFα-treated samples (**Figure 5G**). **Overall, these results show that IL-26 has an immunoregulatory effect under proinflammatory conditions and can decrease the expression of proinflammatory chemokines and cytokines induced by TNFα**.

### IL-26 has a protective role under proinflammatory conditions

Considering the significant changes in the expression of tight junction genes and other tissue-protecting genes, we next performed functional assays to investigate the influence of IL-26 on tissue damage. Within the IO model, epithelial cells form a polarized layer forming the epithelial barrier (**Figure 6A**). TNFα treatment induced tissue damage and impairment of this barrier (**Figure 6B**). This damage was not observed in IL-26-treated samples (**Figure 6C**). More strikingly, IL-26 protected IOs against TNFα-induced tissue damage (**Figure 6D**). To quantify this protective effect, we implemented a wound healing assay using an IO-derived 2D model of intestinal tissue. This analysis revealed delayed wound healing in TNFα-treated, but not IL-26-treated, samples. Remarkably, in the presence of IL-26, the effect of TNFα treatment on wound healing speed was reduced (**Figure 6E**). An Annexin V apoptosis assay subsequently showed a significant decrease of Annexin V^+^ cells in TNFα+IL-26-treated compared to TNFα-treated IOs (**Figure 6F**). **Taken together, these results show that IL-26 plays an important protective role against inflammation-induced intestinal tissue damage.**

**Figure 6.**
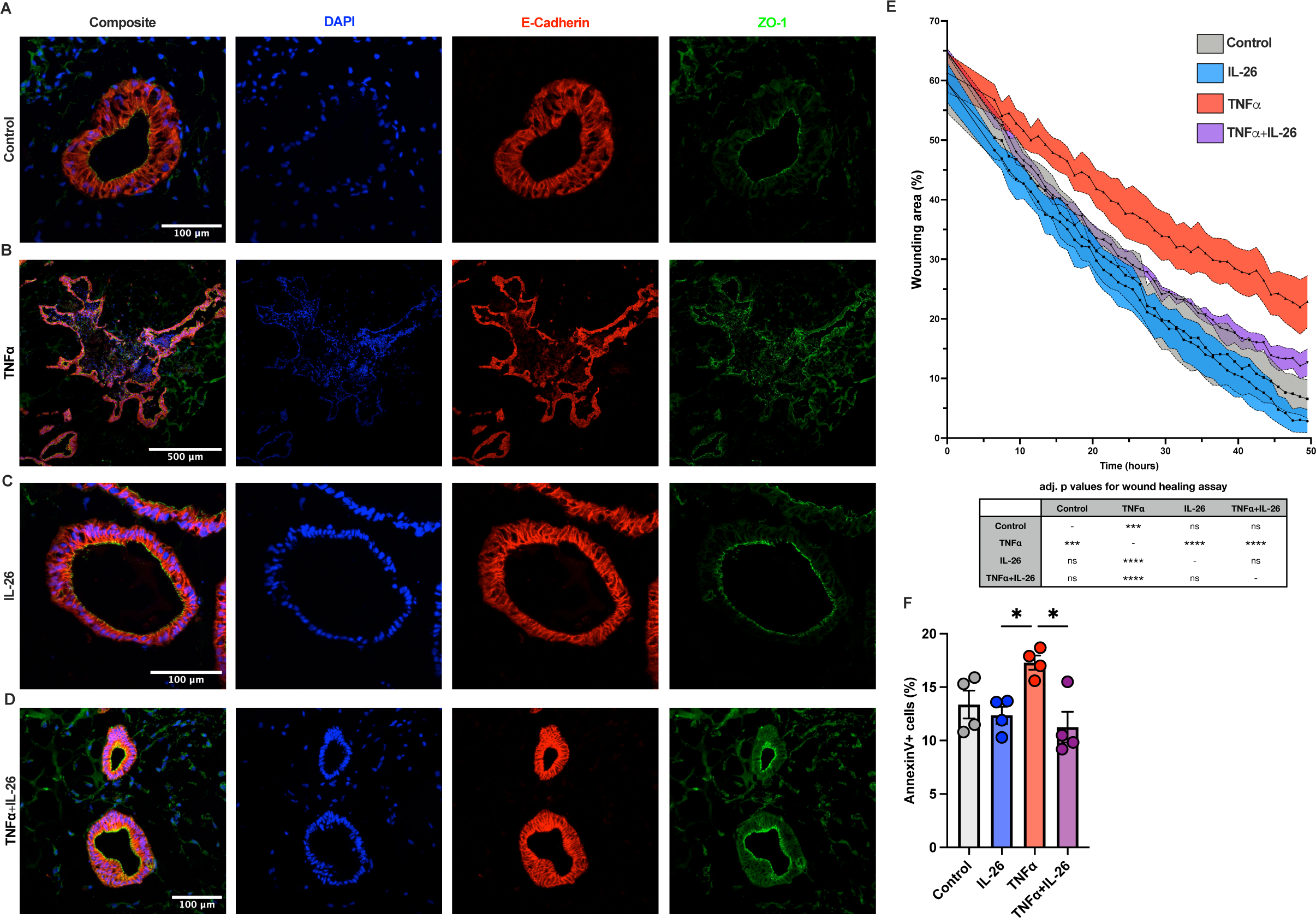
IL-26 protects against inflammation-induced damage in intestinal tissue. (A-D) Representative immunofluorescence images of IOs without treatment (A) or after 24-hour treatment with TNFα (B), IL-26 (C), or TNFα+IL-26 (D). (E) Quantification of wound healing assay performed on an organoid-derived 2D model. Table shows statistical significance of differences between groups. Statistical testing of these differences was performed using the generalized least squares method. (F) Annexin V^+^ cells after 24 hours of stimulation with TNFα, IL-26, or TNFα+IL-26. \**p* < .05, \*\*\**p* < .005, \*\*\*\**p* < .001, ns, not significant.

### Peripheral blood MAIT cells migrate towards IO tissue

We previously showed that IOs do not consist of hematopoietic cells^25^ (**Figure S6A**). However, IOs form an immunocompetent environment^25^ and therefore serve as an ideal model to study the interaction of mucosal tissue with immune cells. Firstly, we used FACS-sorted MAIT cells from HD buffy coats for co-cultivation with IOs. After 24 hours of co-cultivation, whole mount staining (**Figure S6B**) and histological slides of IOs (**Figure S6C**) showed that MAIT cells were in close proximity to IO epithelial cells. Next, to mimic the activated phenotype of MAIT cells in intestinal tissue, we activated FACS-sorted cells with a combination of TCRd and TCRi stimuli and co-cultivated with untreated (**Figure 7A**) and TNFα-treated (**Figure 7B**) IOs. Consistent with our findings for IL-26 (**Figure 6**), we did not observe any TNFα-induced tissue damage in IOs co-cultivated with activated MAIT cells (**Figure 7C, D**). Finally, we used flow cytometric analysis to quantify the percentage of MAIT cells within the IOs, finding that TNFα treatment significantly increased the percentage of CD45^+^ cells in IO tissue (**Figure 7E, F**). **In conclusion, these results show that hiPSC-derived IOs can serve as a model to study interactions between MAIT cells and mucosal tissue. Peripheral blood MAIT cells co-cultivated with IOs can be found in close proximity to epithelial cells, recapitulating the condition *in vivo*, and TNFα treatment increases the percentage of these MAIT cells within the IOs.**

**Figure 7.**
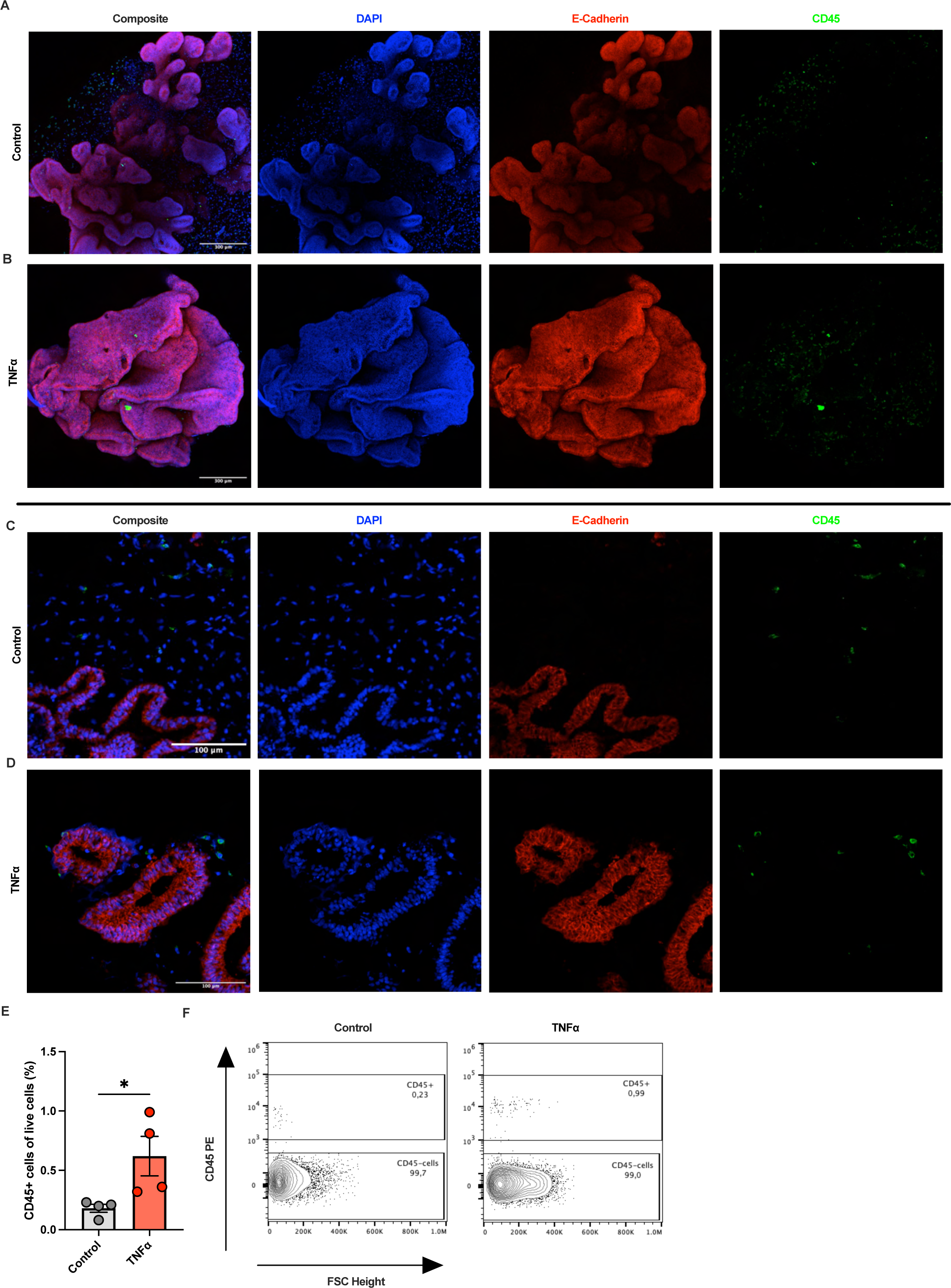
MAIT cells migrate in close proximity to epithelial cells. (A-D) Immunofluorescence staining of whole tissue (A, B) and histological slides (C, D) of IOs. (E) Bar plot showing the percentage of CD45^+^ cells within dissociated IOs co-cultivated with MAIT cells. (F) Representative dot plots showing CD45 expression in IOs co-cultivated with MAIT cells. \**p* < .05.

## DISCUSSION

In this study, we report a decreased percentage and more activated phenotype of MAIT cells during active intestinal inflammation by comparing MAIT cells from non-inflamed and inflamed ileum of CD patients undergoing bowel resection. We compared the expression of markers including CD4 and CD8 in MAIT cells and showed dysregulated expression of IL-26 by these cells in CD patients compared to HD, indicating an important role for IL-26-expressing MAIT cells in intestinal tissue homeostasis. Next, using our state-of-the-art 3D model of human intestinal tissue, we showed the protective role of IL-26 in healthy mucosal tissue and under proinflammatory conditions.

Corroborating previously published findings^11–15^, we found that MAIT cells were significantly decreased in the peripheral blood of CD patients compared to HD. So far, most studies investigating MAIT cells in IBD have focused on peripheral blood, while their role in intestinal tissue during inflammatory conditions remains poorly understood. Serriari *et al.*^13^ used fluorescent microscopy to show tissue accumulation of IL-18Rα^+^TCR Vα7.2^+^CD3^+^ MAIT cells during inflammation. Conversely, Hiejima *et al.*^14^ showed a significant decrease of CD161^high^TCR Vα7.2^+^CD3^+^ MAIT cells in the inflamed tissue of CD patients compared to non-inflamed tissue; however, they did not confirm these results using flow cytometry. In studies of UC, the results have been more consistent, demonstrating a clear accumulation of MAIT cells in the inflamed colon^15, 16^. Our microscopy and flow cytometry results revealed a significant decrease of MAIT cells in inflamed tissue compared to non-inflamed tissue of CD patients. These observations suggest that the role of MAIT cells in IBD is complex and that changes in the MAIT cell population might depend on multiple factors including disease activity, location, or duration. Interestingly, in peripheral blood, the decrease of MAIT cells has been proven as a “general” phenomenon for both CD and UC. In tissue, the changes might be more complicated and affected by more aspects such as diet^35^ or microbiome^36^.

As our results showed an activated phenotype of MAIT cells in inflamed tissue of CD patients, we sought to investigate differences between the physiological changes upon activation of MAIT cells from HD and the pathological changes of activated MAIT cells seen in this inflamed tissue. *In vitro* activation of HD MAIT cells resulted in a decrease of CD8^+^ and increase of DN MAIT cells in culture. As CD4 and CD8 molecules act as TCR co-receptors, they might be crucial for proper and accurate activation of MAIT cells. In our study, CD8^+^ MAIT cells showed increased activation (increased expression of CD69) after *in vitro* TCRi stimulation compared to DN MAIT cells. Moreover, we observed different percentages of CD8^+^ MAIT cells in non-inflamed ileum and peripheral blood of HD, indicating different activation statuses and roles for these cells. Furthermore, we showed that the percentages of CD8^+^ and DN MAIT cells in peripheral blood were different in CD patients compared to HD, suggesting that the function of MAIT cells is impaired in CD patients. These findings conflict with a previous study showing a decrease in CD8^+^ MAIT cells in peripheral blood of CD patients compared to HD; this discrepancy might be explained by the fact that most of the patients included in that previous study were in remission at the time of blood collection^13^. In our study, only patients undergoing surgery were included. On top of the relative numbers of MAIT cells, their phenotypes may also be variable and dynamic during IBD development. Further research is needed to understand correlations and pathophysiological consequences of these phenomena.

Next, we used SCENITH^TM 30^ to compare the cellular energy metabolism of MAIT cells and conventional T cells from peripheral blood of HD. Our results showed a unique energy metabolism profile for MAIT cells, with significantly higher glucose dependence and glycolytic capacity relative to conventional T cells. We hypothesize that glucose levels may be a control mechanism regulating the proper activation of MAIT cells and the production of effector molecules including IL-26. We also observed a significantly higher percentage of activated MAIT cells in peripheral blood of CD patients compared to HD, corroborating previous reports^12, 15^ – this may be attributable to increased glucose levels in blood during active inflammation. Ileum contributes to glucose uptake via digestion of food; therefore, the concentration of glucose can be several times higher in the intestine compared to peripheral blood^37^. Divergent levels of glucose might explain the different phenotypes of MAIT cells in ileum and peripheral blood. Moreover, it has been shown that MAIT cells are involved in the pathology of type 1 (T1D) and 2 (T2D) diabetes – as in IBD, MAIT cells are less abundant in the peripheral blood of T1D^38^ and T2D^39^ patients, and the cells display an activated phenotype^39^. These findings further indicate a link between blood glucose levels and pathological MAIT cell activation.

We and others previously showed the crucial role of TCRi activation in IL-26 expression by MAIT cells^17, 18, 28, 40^. Therefore, we next focused on the expression of IL-26 by MAIT cells. For the first time, we demonstrated IL-26 expression by MAIT cells in peripheral blood and intestinal tissue from CD patients using flow cytometry, which allowed us to quantify IL-26^+^ MAIT cells and thus better understand their involvement during inflammation. We showed that in peripheral blood from HD, MAIT cells did not express IL-26. Conversely, IL-26^+^ MAIT cells were significantly more abundant in peripheral blood from CD patients, in line with previous results showing a higher serum level of IL-26 in CD patients^41^. IL-26 was also highly expressed by MAIT in non-inflamed ileum, indicating its crucial role in the homeostasis of this tissue. We also observed a pathological decrease of IL-26^+^ MAIT cells in inflamed tissue from CD patients. These observations indicate dysregulation of IL-26 expression by MAIT cells in CD patients, both in peripheral blood and ileum, and suggest that MAIT cells and IL-26 may have distinct roles in different tissues and peripheral blood.

Finally, we investigated the role of IL-26 in healthy mucosal tissue by using hiPSC-derived IOs, which form complex organized structures mimicking human intestinal tissue and contain various cell types including epithelial and mesenchymal cells. Previous studies of IL-26 in the context of intestinal inflammation used simple 2D models^41–43^, which lack tissue complexity as they are grown only in monolayer and usually consist of only one cell type. Here, we showed that IL-26 does not induce the expression of proinflammatory chemokines or cytokines in healthy tissue; instead, it induces the expression of genes involved in tight junction assembly and epithelial barrier integrity. Accordingly, we found that IL-26 has a protective role under proinflammatory conditions, significantly reducing TNFα-induced tissue damage. These data corroborate previously published results; Corridoni *et al.*^22^, for instance, showed that IL-26 attenuated acute colitis in a humanized IL-26 murine model. IL-26-expressing mice in that study also exhibited significantly decreased levels of CXCL9 and CXCL10 in inflamed intestinal tissue. We saw a similar pattern here, where the expression of both CXCL9 and CXCL10 in our IO model was increased after TNFα stimulation and reduced in the presence of IL-26.

Previously, we have shown that IOs do not contain any immune cells; however, they form an immunocompetent microenvironment allowing engraftment of organoids with immune cells^25^. Here, we demonstrated that IOs can be used for studying the interaction of MAIT cells with mucosal tissue. Co-cultivation of MAIT cells with TNFα-treated IOs resulted in higher accumulation of MAIT cells in the tissue. These findings can help to better understand the role of MAIT cells in intestinal tissue under homeostatic and inflammatory conditions. This co-cultivation model could be further used to investigate possible targets to inhibit pathological activation of MAIT cells or their stimulation in order to increase IL-26^+^ cells in inflamed tissue.

Overall, our study provides important insight into the biology of MAIT cells and IL-26 and their roles in the maintenance of intestinal homeostasis and IBD. For the first time, we have shown expression of IL-26 by MAIT cells using flow cytometry and described its dysregulation in CD patients. We combined findings from a clinical cohort with state-of-the-art *in vitro* methods to characterize our observations. Using single-cell energy metabolism analysis (SCENITH^TM^), we have shown that MAIT cells represent a distinct population of T cells with higher glucose dependency. Using a complex 3D human *in vitro* model, we have shown a clear protective role of IL-26 in healthy and inflamed intestinal tissue. These findings might pave the road for the development of new therapeutics targeting MAIT cells and IL-26. Overall, our data suggest that restoring IL-26 secretion from MAIT cells may have a protective effect on the epithelial barrier, potentially aiding in recovery during IBD flares. However, further analysis of the molecular pathways underlying the protective role of IL-26 and of how the modulation of MAIT cell metabolism could be used to treat CD patients is needed. Furthermore, our study focused only on ileum of CD patients, therefore further research is needed to show involvement of MAIT cells and IL-26 in other parts of gastrointestinal tract such as colon.

### Conflict of interests

**B Verstockt reports:** Research support from AbbVie, Biora Therapeutics, Landos, Pfizer, Sossei Heptares and Takeda. Speaker’s fees from Abbvie, Biogen, Bristol Myers Squibb, Celltrion, Chiesi, Falk, Ferring, Galapagos, Janssen, Lily, MSD, Pfizer, R-Biopharm, Sandoz, Takeda, Tillots Pharma, Truvion and Viatris. Consultancy fees from Abbvie, Alfasigma, Alimentiv, Applied Strategic, Astrazeneca, Atheneum, BenevolentAI, Biora Therapeutics, Boxer Capital, Bristol Myers Squibb, Galapagos, Guidepont, Landos, Lily, Merck, Mylan, Inotrem, Ipsos, Janssen, Pfizer, Progenity, Sandoz, Sanofi, Santa Ana Bio, Sapphire Therapeutics, Sosei Heptares, Takeda, Tillots Pharma and Viatris. Stock options Vagustim. **S Vermeire reports:** Grants from AbbVie J&J Pfizer Galapagos and Takeda consulting and/or speaking fees from AbbVie Abivax AbolerIS Pharma AgomAb Alimentiv Arena Pharmaceuticals AstraZeneca Avaxia BMS Boehringer Ingelheim Celgene CVasThera Dr Falk Pharma Ferring Galapagos Genentech-Roche Gilead GSK Hospira Imidomics Janssen J&J Lilly Materia Prima MiroBio Morphic MrMHealthMundipharma MSD Pfizer Prodigest Progenity Prometheus Robarts Clinical Trials Second Genome Shire Surrozen Takeda Theravance Tillots Pharma AG and Zealand Pharma.

All other authors declare no conflicting interest.

### Author contributions

Conceptualization: VB, BJK, MHK, MDZ, GM, JF; Methodology: VB, BJK, MHK, MDZ, IDG, SaV, BV, SeV, RJA, GM, JF; Investigation: VB, BJK, FK, IP, SS, FB, SA; Visualization: VB, IP, BJK; Funding acquisition: BV, SeV, GM, JF; Supervision: GM, JF; Writing -original draft: VB; Writing - review and editing: all.

## ACKNOWLEDGMENTS

We would like to thank all patients for their participation in the clinical study and for their willingness to donate tissue samples for research. We thank the IBD Leuven group for assistance with patient samples collection and administration. We thank the technical support team of the Center for Translational Medicine for technical assistance. We acknowledge the CF Genomics CEITEC MU supported by the NCMG research infrastructure (LM2023067 funded by MEYS CR) for their support with obtaining scientific data presented in this paper. Core Facility Bioinformatics of CEITEC Masaryk University is gratefully acknowledged for the obtaining and analyzing of the scientific data (RNA-seq) presented in this paper. The authors gratefully acknowledge the Biostatistics Core Facility of FNUSA-ICRC for their support and assistance in this work. We thank Tobie Martens (TARGID, KU Leuven) for his assistance and support with Zeiss LSM 780, Reena Chinnaraj and Vera Dermesrobian (FACS Core, KU Leuven) for support with flow cytometry. Graphical images were generated using BioRender. We would like to thank Dr. Daniel Ackerman from Insight Editing London for critical review and editing of the manuscript prior to submission.

## Abbreviations used in this paper

**Main text**
3D: 3-dimensional
CD: Crohn’s disease
CD8^+^: CD4^-^CD8^+^
DEGs: differentially expressed genes
DG: 2-deoxy-D-glucose
DN: double-negative (CD4^-^CD8^-^)
FACS: fluorescence-activated cell sorting
FAO/AAO: fatty and amino acid oxidation
HD: healthy donor
hiPSC: human induced pluripotent stem cell
hrIL-26: human recombinant IL-26
IBD: inflammatory bowel disease
INF: inflamed
IOs: intestinal organoids
MAIT cells: mucosal-associated invariant T cells
MR1: MHC class I-related molecule 1
NON-INF: non-inflamed
O: oligomycin
PBMCs: peripheral blood mononuclear cells
RNA-seq: RNA sequencing
T1D: type 1 diabetes
T2D: type 2 diabetes
TCR: T-cell receptor
TCRd: TCR-dependent
TCRi: TCR-independent
UC: ulcerative colitis

**Material and Methods**
2D: 2-dimensional
BSA: bovine serum albumin
DE: definitive endoderm
FBS: fetal bovine serum
HBSS: Hank’s balanced salt solution
PBS: phosphate-buffered saline
RT: room temperature

## Notes

**Grant support:**

This study was supported by fundings - European Union - Next Generation EU project nr. LX22NPO5107, European Regional Development Fund - Project (ENOCH, CZ.02.1.01/0.0/0.0/16_19/0000868), Ministry of Health of the Czech Republic-DRO (Institute of Hematology and Blood Transfusion-00023736). This study was supported by the Support of Internal Pilot Research Projects of the St. Anne’s University Hospital in Brno. G.M.’s lab was supported by FWO grants G0D8317N, G0A7919N, G086721N, G088816N and S008419N (SV and GM), KU Leuven Internal Funds (C12/15/016 and C14/17/097 (SV and GM)), European Union’s Horizon Europe Research & Innovation programme (FIBROTARGET 101080523, BJ, SV and GM) and European Crohn’s and Colitis Organisation (ECCO) Pioneer Award 2023 (GM), Helmsley Charitable Trust (SV and GM), Taiwan (Ministry of Education)-KU Leuven Scholarship Programme (BJK). BV is supported by the Clinical Research Fund (KOOR) at the University Hospitals Leuven and the Research Council at the KU Leuven. SV holds a BOF-FKO from the KU Leuven.

### Competing Interest Statement

B Verstockt reports: Research support from AbbVie, Biora Therapeutics, Landos, Pfizer, Sossei Heptares and Takeda. Speaker fees from Abbvie, Biogen, Bristol Myers Squibb, Celltrion, Chiesi, Falk, Ferring, Galapagos, Janssen, Lily, MSD, Pfizer, R-Biopharm, Sandoz, Takeda, Tillots Pharma, Truvion and Viatris. Consultancy fees from Abbvie, Alfasigma, Alimentiv, Applied Strategic, Astrazeneca, Atheneum, BenevolentAI, Biora Therapeutics, Boxer Capital, Bristol Myers Squibb, Galapagos, Guidepont, Landos, Lily, Merck, Mylan, Inotrem, Ipsos, Janssen, Pfizer, Progenity, Sandoz, Sanofi, Santa Ana Bio, Sapphire Therapeutics, Sosei Heptares, Takeda, Tillots Pharma and Viatris. Stock options Vagustim. S Vermeire reports: Grants from AbbVie J&J Pfizer Galapagos and Takeda consulting and/or speaking fees from AbbVie Abivax AbolerIS Pharma AgomAb Alimentiv Arena Pharmaceuticals AstraZeneca Avaxia BMS Boehringer Ingelheim Celgene CVasThera Dr Falk Pharma Ferring Galapagos Genentech-Roche Gilead GSK Hospira Imidomics Janssen J&J Lilly Materia Prima MiroBio Morphic MrMHealthMundipharma MSD Pfizer Prodigest Progenity Prometheus Robarts Clinical Trials Second Genome Shire Surrozen Takeda Theravance Tillots Pharma AG and Zealand Pharma.
All other authors declare no conflicting interest.

