## Supplementary information for "Unveiling the guardians: IL-26-expressing MAIT cells protect epithelial barrier function and are dysregulated in Crohn’s disease"

This Supplementary information contains Supplementary Materials and Methods, Figure S1, Figure S2, Figure S3, Figure S4, Figure S5, Figure S6. Table S1, Table S2, and Table S3 showing details of clinical data of patients included in the study, used antibodies and TaqMan$^{TM}$ probes.

### SUPPLEMENTARY MATERIALS AND METHODS

#### Immunofluorescence staining of tissue samples

Tissue was fixed with 4% formaldehyde and embedded in paraffin as the standard protocol at the UZ Leuven. The samples were cut into 5 µm sections. The samples were deparaffinized and rehydrated and the antigens were retrieved using sodium citrate buffer (pH 6) for 20 minutes at 95 °C. The sections were permeabilized using 0.2% Triton X-100 in phosphate-buffered saline (PBS) with 0.1% Tween 20 (PBST) for 10 minutes, and unspecific binding sites were blocked with 2% bovine serum albumin (BSA) in PBST for 60 minutes. Samples were incubated with primary antibodies anti-human TCR Vα7.2, IL-26, KLRB1 in PBST + 2% BSA overnight at 4 °C. After washing, the samples were labeled with secondary antibodies (all antibody details are shown in **Table S2**) in PBST + 2% BSA for 2 hours at room temperature (RT). Before data acquisition, the cells were counterstained with DAPI for 15 minutes at RT. MAIT cells were counted as $CD161^{+}$TCR $V\alpha7.2^{+}$ cells as three technical replicates for three individual patients.

#### Isolation of single-cell suspension from the ileum

Cells were isolated from the transmural terminal ileum of CD patients undergoing surgery. In brief, the tissue was incubated with 1 mM dithiothreitol (DTT) and 1 mM EDTA in Hank's balanced salt solution (HBSS) supplemented with 1% fetal bovine serum (FBS) and 1% Penicillin/Streptomycin (P/S) for 30 minutes at 37 °C. The tissue was treated with 1 mM EDTA in HBSS supplemented with 1% FBS and 1% P/S for 30 minutes at 37 °C. Next, the tissue was cut with sterile scissors and digested with 5.4 U/mL collagenase D (Roche Applied Science), 39.6 U/mL dispase II (Gibco) and 100 U/mL DNAse I (Sigma) in a gentleMACS C tube (Miltenyi Biotec) for 20 minutes at 37 °C after dissociation with the gentleMACS Dissociator (program human_tumor_02.01). The cell suspension was treated with Red Blood Cell lysis buffer (Roche). The cells were used for flow cytometry.

#### Peripheral blood mononuclear cells (PBMCs) isolation

PBMCs were isolated either from whole blood samples (the blood was collected in the EDTA tube from healthy donors or CD patients one day before surgery, the protocol was approved by the Institutional Review Board (IRB) of the University Hospitals Leuven, Belgium (B322201213950/S53684, CCARE, S-53684)) or fresh healthy buffy coats (Department of Transfusion & Tissue Medicine of the Brno University hospital). The blood samples were diluted with 2% FBS in PBS prior to gradient centrifugation on Lymphoprep (STEMCELL Technologies).

**MAIT cells isolation**

Fluorescence-activated Cell Sorting (FACS) was used for MAIT cell isolation. Briefly, the PBMCs isolated from fresh healthy buffy coats were stained with Streptavidin and fluorochrome-conjugated anti-human CD3, CD14, CD19, CD161, TCR Vα7.2 antibodies (clones and dilutions are shown in **Table S2**). Dead cells were excluded using LIVE/DEAD™ Fixable Near-IR Dead Cell Stain Kit (Invitrogen). The cells were sorted as Live/dead$^{-}$Lin(CD14, CD19)$^{-}$CD3$^{+}$CD161$^{+}$TCR Vα7.2$^{+}$ using a MoFlo Astrios EQ Cell Sorter (Beckman Coulter).

***In vitro* activation of MAIT cells**

For *in vitro* activation, isolated PBMCs were stimulated for 48 hours with cytokines (TCRi) including IL-12 (2 ng/mL, R&D), IL-15 (25 ng/mL, R&D), IL-18 (50 ng/mL, R&D), and TL-1A (100 ng/mL, R&D), or with anti-CD3 and -CD28 beads (TCRd) from T Cell Activation/Expansion Kit (Miltenyi Biotec), or in combination (as defined in **Figure 3B**).

**Intracellular staining for *in vitro* expression of IL-26**

After stimulation, the cells were stained for surface markers using fluorochrome-conjugated anti-human CD3, CD4, CD8, CD161, TCR Vα7.2 antibodies. After staining, the cells were fixed and permeabilized using the eBioscience IC staining kit. The cells were stained with anti-human IL-26 antibody (all antibody details are shown in **Table S2**). The dead cells were excluded using LIVE/DEAD™ Fixable Violet Dead Cell Stain Kit (Invitrogen). The MAIT cells were identified as the Live/dead$^{-}$CD3$^{+}$CD161$^{+}$TCR Vα7.2$^{+}$ cell population. The samples were analyzed on an SA3800 spectral flow cytometer (Sony Biotechnology).

**Annexin V staining**

FACS-sorted MAIT cells were stimulated (according to the aforementioned protocol) for 72 hours. Prior to the Annexin V staining, the cells were stained with anti-human CD4, CD8, CD45, CD69, CD161, and TCR Vα7.2 antibodies (details in **Table S2**). For the detection of apoptotic cells, the Annexin V Apoptosis Detection Kit (eBioscience) was used. Briefly, the cells were washed with PBS and a binding buffer. Next, the cells were incubated with eFluor450-conjugated Annexin V for 15 minutes at RT. Dead cells were excluded using 7-AAD Viability Staining Solution (eBioscience). The samples were analyzed on an SA3800 spectral flow cytometer (Sony Biotechnology).

**Single-cell energy metabolism profiling (SCENITH™)**

PBMCs isolated from fresh healthy buffy coats were used for SCENITH™. After isolation, the cells were rested for 1 hour at 37 °C in RPMI-1640 medium (Gibco) supplemented with 2% FBS and 1% P/S. For *in vitro* activation, the cells were stimulated with TCRi and TCRd stimuli according to the aforementioned protocol for 2 hours prior to SCENITH™. The analysis was performed according to a previously published protocol[30]. The cells were stained with anti-human CD3, CD4, CD8, CD14, CD19, CD161, and TCR Vα7.2 antibodies and Streptavidin (details in **Table S2**). The dead cells were excluded using LIVE/DEAD™ Fixable Green Dead Cell Stain Kit (Invitrogen). The MAIT cells were identified as Live/dead$^-$Lin(CD14, CD19)$^-$ CD3$^+$CD161$^+$TCR Vα7.2$^+$. The samples were analyzed on an SA3800 spectral flow cytometer (Sony Biotechnology).

**RNA extraction and gene expression analysis**

The cells were lysed in TRI Reagent® (Merck) and the RNA was isolated by column separation using RNeasy Plus Micro Kit with gDNA elimination (Qiagen) according to the manufacturer's protocol. Gene expression was analyzed using qPCR TaqMan probes listed in **Table S3** and TaqMan™ Gene expression Master Mix (Thermo Fisher Scientific) according to the manufacturer's protocol. For the qPCR analysis, the StepOne™ Real-Time PCR System (Applied Biosystems) was used.

**Human induced pluripotent stem cells (hiPSCs) maintenance**

hiPSCs (WiCell, DF19-9-7T[44]) were cultured on Matrigel-coated tissue culture dishes and were maintained in mTeSR Plus medium (STEMCELL Technologies). The medium was changed every two days. When the cells reached ~80% confluency, they were passaged using TryPLE (Gibco), and the RHO/ROCK pathway inhibitor (STEMCELL Technologies) was used for 24 hours.

**Dissociation of IOs**

IOs were dissociated to obtain single cell suspension. Briefly, the organoids were washed with cold HBSS and mechanically disrupted with sterile scissors. The cells were dissociated with TryPLE for 10 minutes at 37 °C. After digestion, the cells were filtered using a 70 µm strainer and washed with cold HBSS + 2% FBS. The single cell suspension was used for further experiments including IO-derived 2D models, and flow cytometry.

**Immunofluorescence staining for IOs**

IOs were washed with cold PBS and fixed with 4% paraformaldehyde for 20 minutes at RT. For whole mount staining, the organoids were permeabilized according to following protocol. For histological slides, the IOs were firstly dehydrated using 15% sucrose overnight at 4 °C. Next, the samples were frozen in Tissue Freezing Medium (Leica Biosystems). The frozen samples were cut and rehydrated in PBS for 10 minutes. The cells were permeabilized with PBS + 0.5% Triton-X100 for 15 minutes at RT and washed with IF buffer (PBS + 0.2% Triton-X100 + 0.05% Tween 20). Samples were blocked with IF buffer + 2.5% BSA for one hour at RT. IOs were labeled with primary anti-human ZO-1, E-Cadherin, IL-10Rβ, IL-20Rα, and CD90 antibodies overnight at 4 °C, then washed three times with IF buffer and labeled with secondary antibodies (all antibody details are shown in **Table S2**) in IF buffer with 1% BSA for 2 hours at RT. IOs were washed three times with IF buffer and incubated with DAPI for 5 minutes at RT. Sections were imaged with a Zeiss LSM 780 confocal microscope using 10x magnification. The images were processed using ImageJ software.

**Stimulation of the IOs**

Matured IOs were stimulated with 50 ng/mL IL-26 (R&D), 10 ng/mL TNFα (R&D), or a combination of both in complete IOs medium. For qPCR and RNA-seq experiments, IOs were stimulated for 4 hours. For the bead array cytokines detection, IOs were stimulated for 24 hours. For the western blot, IOs were stimulated with IL-26 for 10, 30, or 60 minutes.

**Bead array cytokines detection**

Matured IOs were stimulated with IL-26 (as described above). For cytokine production measurement, the LEGENDplex™ Human Inflammation Panel 13-plex cytometric bead array (BioLegend) was used according to the manufacturer's protocol.

**RNA sequencing (RNA-seq)**

IOs stimulated with IL-26, TNFα, or TNFα+IL-26 and nonstimulated controls (as described above) were washed with cold PBS and the Cultrex was removed using Corning® Cell Recovery Solution (Corning) for 20 minutes at 4 °C. Cells were lysed in of TRI Reagent®. The RNA was isolated using the RNeasy mini kit (Qiagen). The integrity of isolated RNA was measured with the Bioanalyzer2100 RNA nano 6000 chips (Agilent Technologies). All analyzed samples demonstrated RIN ≥ 9.9. Libraries were constructed with the QuantSeq 3' mRNA-Seq Library Prep Kit FWD using 250 ng of RNA. Samples were sequenced using the Lexogen QuantSeq FWD kit at a sequencing depth of ~13M for all samples. Raw reads were quality checked (FastQC, MultiQC, minion, swan), preprocessed (Trimmomatic) and mapped (STAR, Samtools) to the reference genome with gene annotation (genome version: Ensembl GRCh38, gene annotation: Ensembl v94). Differential expression (DE) analysis was done in the R v4.3.1 environment using the DESeq2 pipeline. The DE analysis results were filtered for low counts and genes were considered differentially expressed (DEGs) when they demonstrated a $|\log2(\text{FoldChange})| \geq 0.6$. Gene ontology (GO) and gene set enrichment analysis (GSEA) were performed with the package clusterProfiler in R or with the desktop version of GSEA MSigDB respectively. The visualization of the results was performed with the R package ggplot2.

**Western blot (WB)**

IOs were washed with cold PBS and lysed with RIPA buffer with Halt™ Protease and Phosphatase Inhibitor Cocktail (Thermo Scientific). Lysates were cleared from precipitates by centrifugation at 10,000 g for 10 minutes. The protein lysates were used for immunoblotting with anti-human phospho (Tyr705) STAT3, STAT3, IL-10Rβ, and IL-20Rα antibodies followed by incubation with secondary antibodies. Anti-GAPDH antibody (all antibody details are shown in **Table S2**) was used as an internal control.

**Wound healing assay**

The single-cell suspension from IOs obtained using the aforementioned protocol was seeded on Corning Matrigel Matrix (Corning)-coated plates. The cells were incubated for 14 days to reach confluency prior to the experiments. During the assay, the wound was made mechanically with a pipette tip. Next, the cells were washed and stimulated with IL-26, TNFα, or TNFα+IL-26. The cells were imaged with a Zeiss LSM 780 confocal microscope using 10x magnification and bright-field live imaging for 48 hours. The quantification of the assay was done in ImageJ.

**Apoptosis assay for IOs**

The IOs were stimulated with IL-26, TNFα, or TNFα+IL-26 for 24 hours prior to the assay. The IOs were dissociated according to the aforementioned protocol, and the single-cell suspension was used for the detection of apoptotic cells using the Annexin V Apoptosis Detection Kit as described above. The samples were analyzed on an SA3800 spectral flow cytometer (Sony Biotechnology). Data were analyzed using FlowJo v10 software (BD Life Sciences).

**Co-cultivation of MAIT cells with IOs**

FACS-sorted MAIT cells were stimulated according to the aforementioned protocol for 48 hours. Subsequently, MAIT cells were washed with PBS. Matured IOs were intensively washed with cold PBS. MAIT cells were mixed with IOs ($10^5$ MAIT cells per organoid) and seeded in a drop of Cultrex Membrane Extract, Type 2. The cells were cultivated in IO medium without growth factors. For experiments with TNFα, 10 ng/mL TNFα was added to the medium for 24 hours. MAIT cells were cultivated with IOs for 24 hours. The cells were used for immunofluorescence staining or dissociated for flow cytometric analysis according to the aforementioned protocols. For immunofluorescence staining, anti-human E-Cadherin, CD45 and CD90 antibodies were used. For cytometric analysis, 7-AAD Viability Staining Solution was used to exclude death cells, and anti-human CD45, EpCam antibodies were used for staining (all antibody details are shown in **Table S2**).

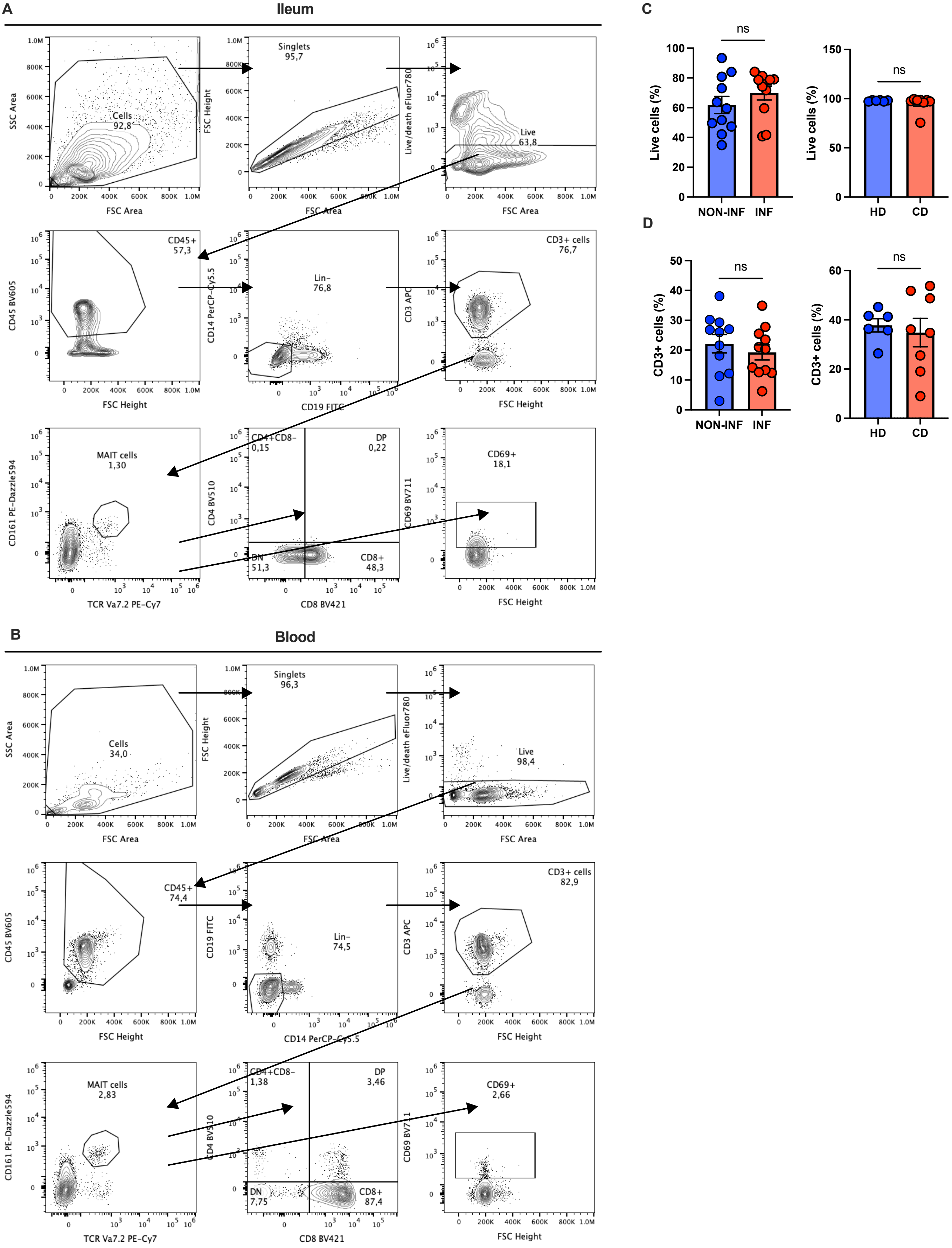

A
Ileum
SSC Area
FSC Area
Cells
92,8
FSC Height
Singlets
95,7
Live/death eFluor780
Live
63,8
CD45 BV605
CD45+
57,3
CD14 PerCP-Cy5.5
Lin-
76,8
CD19 FITC
CD3 APC
CD3+ cells
76,7
FSC Height
CD161 PE-Dazzle594
MAIT cells
1,30
TCR Va7.2 PE-Cy7
CD4 BV510
CD4+CD8-
0,15
DP
0,22
DN
51,3
CD8+
48,3
CD8 BV421
CD69 BV711
CD69+
18,1
B
Blood
Cells
34,0
Singlets
96,3
Live
98,4
CD45+
74,4
CD19 FITC
Lin-
74,5
CD14 PerCP-Cy5.5
CD3+ cells
82,9
MAIT cells
2,83
CD4+CD8-
1,38
DP
3,46
DN
7,75
CD8+
87,4
CD69+
2,66
C
Live cells (%)
ns
NON-INF
INF
HD
CD
D
CD3+ cells (%)
ns
NON-INF
INF
HD
CD

**Figure S1. Identification of MAIT cells in peripheral blood and intestinal tissue.**
(A, B) Gating strategy for flow cytometric analysis of MAIT cells in intestinal tissue (A) and peripheral blood (B). (C, D) Flow cytometric analysis showing the percentages of live cells (C) and $CD3^{+}$ cells (D) among different samples. Statistical significance was calculated using paired *t*-test (for tissue samples) or unpaired *t*-test (for blood samples). ns, not significant.

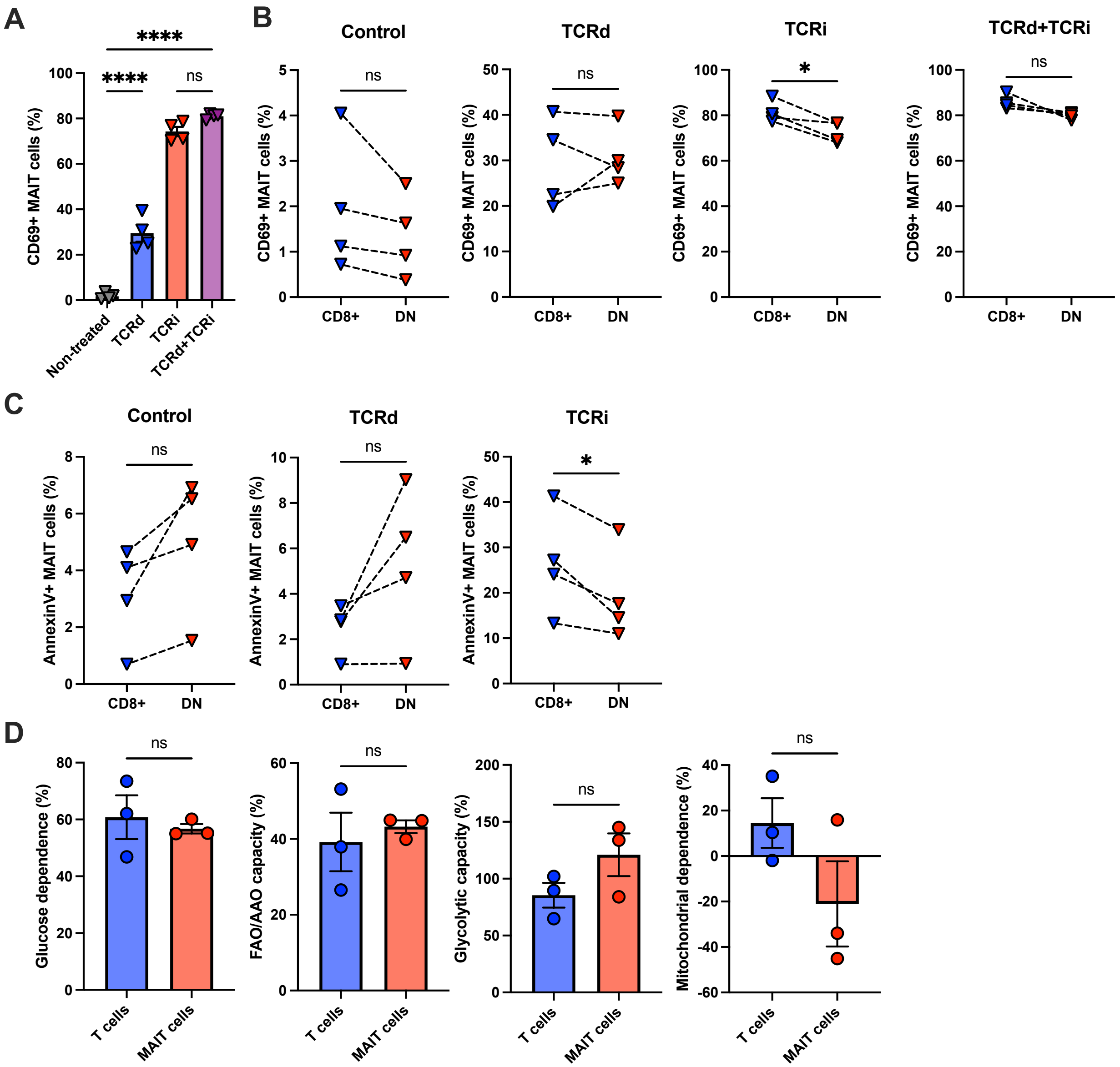

A
****
****
ns
CD69+ MAIT cells (%)
Non-treated
TCRd
TCRi
TCRd+TCRi
B
Control
TCRd
TCRi
TCRd+TCRi
ns
ns
*
ns
CD8+
DN
C
Control
TCRd
TCRi
ns
ns
*
AnnexinV+ MAIT cells (%)
CD8+
DN
D
Glucose dependence (%)
FAO/AAO capacity (%)
Glycolytic capacity (%)
Mitochondrial dependence (%)
ns
T cells
MAIT cells

**Figure S2.** ***In vitro*** **activation of peripheral blood MAIT cells.** (A) Bar plot showing the percentage of $CD69^+$ MAIT cells after *in vitro* stimulation. (B) Percentage of $CD69^+$ MAIT cells at steady state and after TCRd, TCRi, and TCRd+TCRi activation. (C) Percentage of Annexin $V^+$ MAIT cells at steady state and after TCRd and TCRi stimulation. (D) Bar plots showing glucose dependence (left), FAO/AAO capacity (left middle), glycolytic capacity (right middle), and mitochondrial dependence (right) of MAIT cells and T cells after TCRd+TCRi activation. Dashed lines indicate paired data points. *$p < .05$, ****$p < .001$.

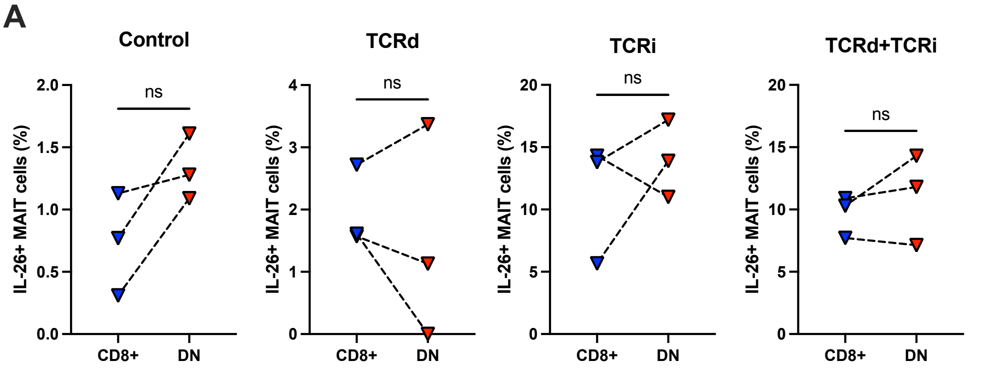

A
Control
TCRd
TCRi
TCRd+TCRi
IL-26+ MAIT cells (%)
ns
CD8+
DN
2.0
1.5
1.0
0.5
0.0
4
3
2
1
0
20
15
10
5
0

**Figure S3. IL-26 Expression by CD8$^+$ and DN MAIT cells.**

(A) Percentage of IL-26$^+$ MAIT cells at steady state and after TCRd, TCRi and TCRd+TCRi activation. Dashed lines indicate paired data points. ns, not significant.

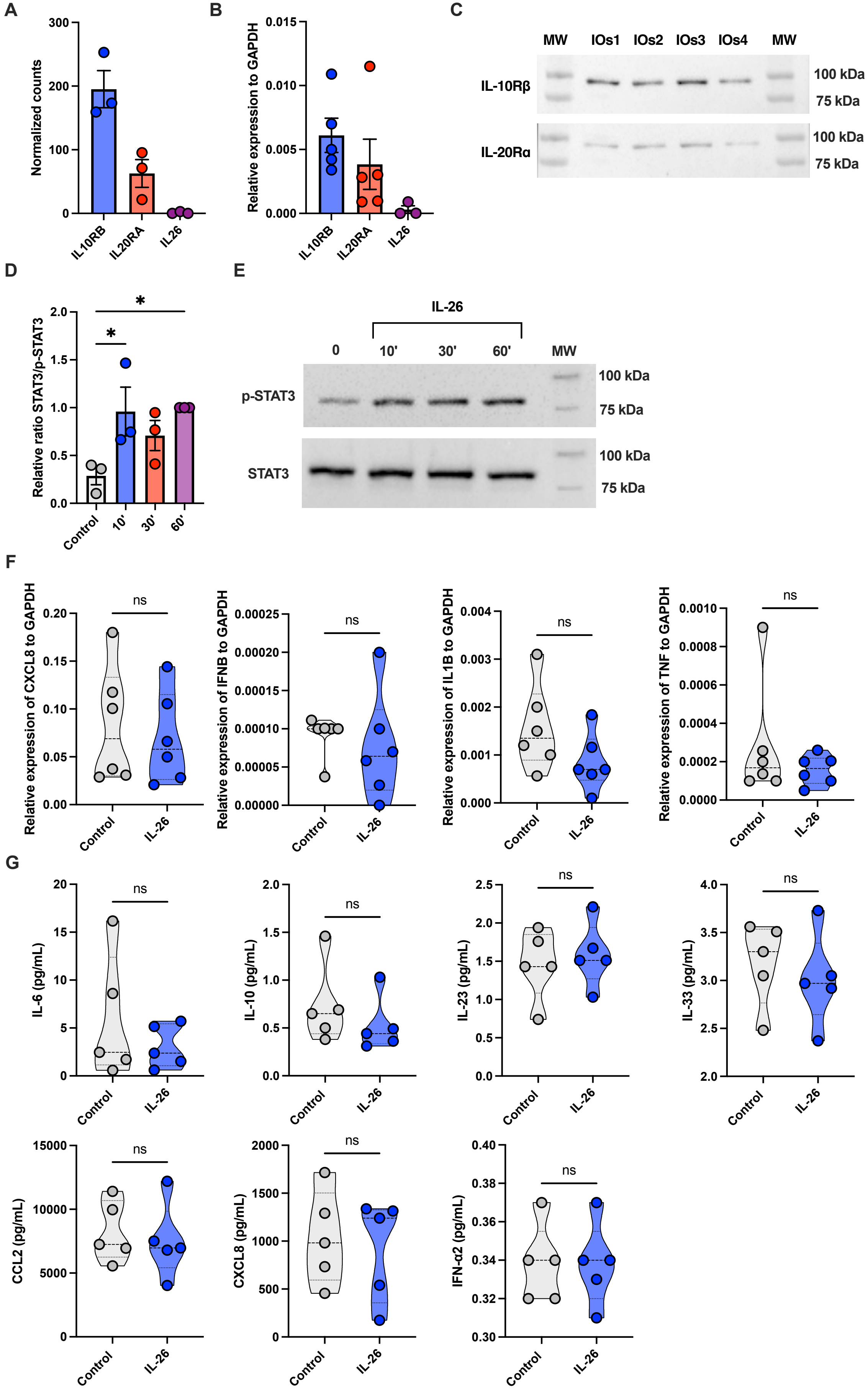
A
Normalized counts
IL10RB
IL20RA
IL26
B
Relative expression to GAPDH
C
MW IOs1 IOs2 IOs3 IOs4 MW
IL-10Rβ
IL-20Rα
100 kDa
75 kDa
D
Relative ratio STAT3/p-STAT3
Control
10'
30'
60'
E
IL-26
0
p-STAT3
STAT3
F
Relative expression of CXCL8 to GAPDH
Relative expression of IFNB to GAPDH
Relative expression of IL1B to GAPDH
Relative expression of TNF to GAPDH
ns
G
IL-6 (pg/mL)
IL-10 (pg/mL)
IL-23 (pg/mL)
IL-33 (pg/mL)
CCL2 (pg/mL)
CXCL8 (pg/mL)
IFN-α2 (pg/mL)

**Figure S4. IL-26 does not induce expression of proinflammatory cytokines and chemokines in healthy intestinal tissue.**

(A-C) IL-26 expression and its receptor in intestinal organoids on the RNA (A, B) and protein (C) levels. (D) Bar plot showing phosphorylation of STAT3 in IL-26 stimulated organoids. (E) Representative western blot analysis of IL-26-stimulated organoid tissue. (F, G) Violin plots showing qPCR analysis (F) and bead array cytokines detection (G) of IL-26-stimulated organoids, dashed lines represent median and dotted lines represent quartiles. *$p$ < .05.

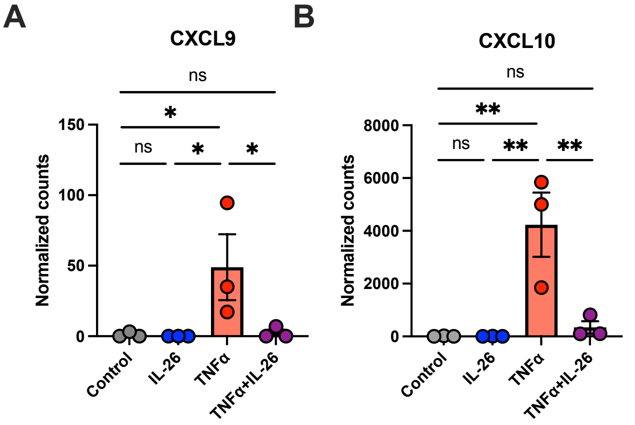

A
CXCL9
ns
*
ns
*
*
Normalized counts
150
100
50
0
Control
IL-26
TNFα
TNFα+IL-26
B
CXCL10
ns
**
ns
**
**
Normalized counts
8000
6000
4000
2000
0
Control
IL-26
TNFα
TNFα+IL-26

**Figure S5. IL-26 decreases expression of proinflammatory chemokines induced by TNFα.**
(A) Bar plots showing expression of CXCL9 and CXCL10 in IOs stimulated with TNFα, IL-26, or TNFα+IL-26. *$p < .05$, **$p < .01$.

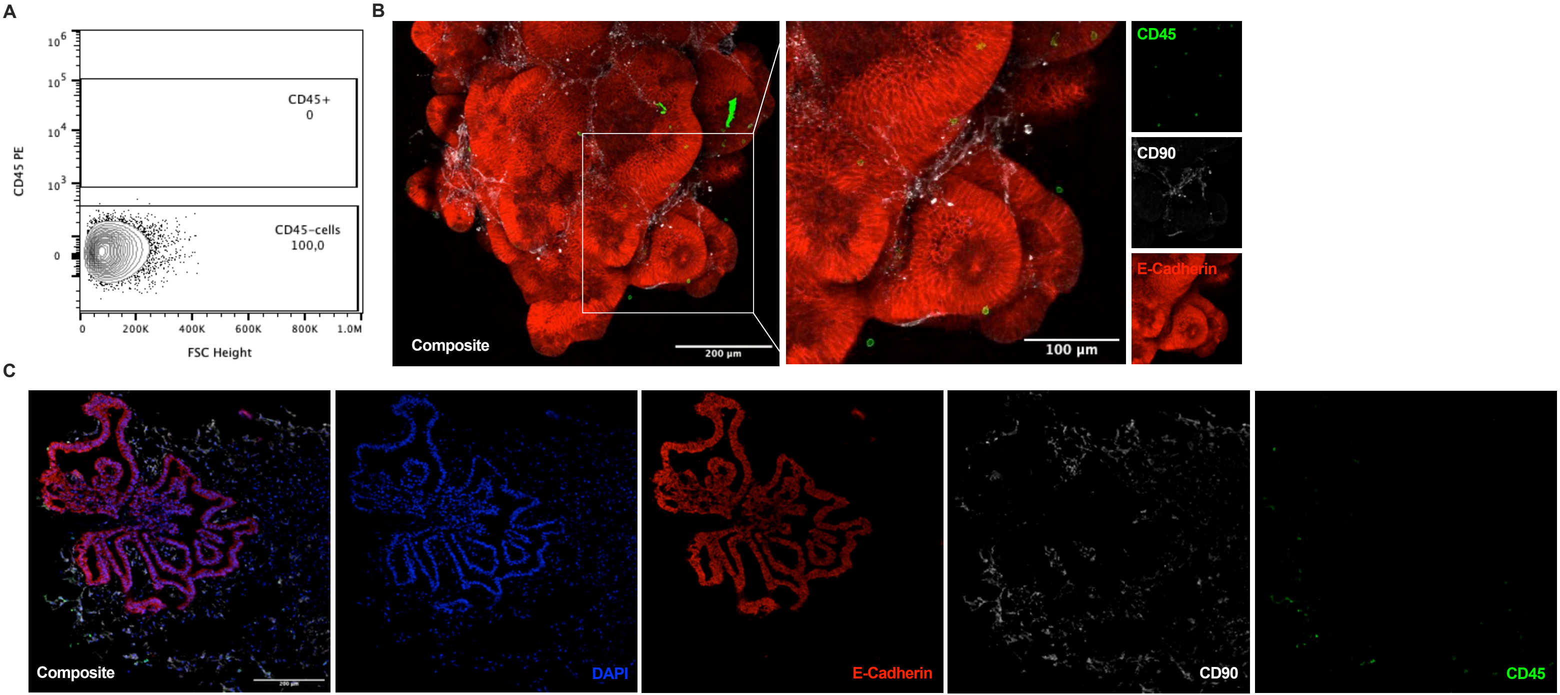

A
B
C
CD45 PE
CD45+
0
CD45-cells
100,0
FSC Height
0
200K
400K
600K
800K
1.0M
Composite
200 µm
100 µm
CD45
CD90
E-Cadherin
DAPI

**Figure S6. Peripheral blood MAIT cells interact with IO tissue.**
(A) Representative dot plot showing no expression of CD45 in IOs. (B, C) Immunofluorescence staining of whole IO tissue (B) and histological slides of IOs (C).

**Table S1 Patients data**

| Characteristic | CD patients (N=16) | Healthy donors (N=6) |
|---|---|---|
| General data | | |
| Age (y/o; mean±SD) | 33.4±14.5 | 27.7±1.4 |
| Sex (M/F) | 11/5 | 2/4 |
| Sample type | | |
| Blood | 8 | 6 |
| Tissue | 11 | 0 |
| Montreal classification | | |
| Location | | |
| Ileal (L1) | 8 | NA |
| Colonic (L2) | 0 | NA |
| Ileocolonic (L3) | 8 | NA |
| Isolated upper disease (L4) | 1 | NA |
| Behaviour | | |
| Non-stricturing, non-penetrating (B1) | 0 | NA |
| Stricturing (B2) | 10 | NA |
| Penetrating (B3) | 6 | NA |
| Perianal disease modifier (P) | 4 | NA |
| Medication at time of surgery | | |
| Medication naive | 2 | NA |
| anti-TNFα (infliximab, adalimumab) | 6 | NA |
| Antibiotics (ciprofloxacin, amoxicillin-clavulanic acid, metronidazole) | 4 | NA |
| Vedolizumab | 3 | NA |
| Steroids (methylprednison, budesonide) | 2 | NA |
| Immunomodulators (Ozanimod, Azathioprin) | 2 | NA |
| Upadacitinib | 1 | NA |
| Risankizumab | 1 | NA |

**Table S2 Table of used antibodies with clones and dilutions**

| Antibody | Conjugate | Clone | Dilution | Vendor | Host |
|---|---|---|---|---|---|
| CD14 | PerCP/Cy5.5 | HCD14 | 1:300 | BioLegend | Mouse |
| CD14 | Biotin | 61D3 | 1:100 | Invitrogen | Mouse |
| CD161 (KLRB1) | PE/Dazzle594 | HP-3G10 | 1:50 | Sony | Mouse |
| CD161 (KLRB1) | - | Polyclonal | 1:50 | Sigma-Aldrich | Rabbit |
| CD19 | FITC | HIB19 | 1:300 | Invitrogen | Mouse |
| CD19 | Biotin | HIB19 | 1:100 | Invitrogen | Mouse |
| CD3 | APC | SK7 | 1:100 | Sony | Mouse |
| CD3 | BV650 | UCHT | 1:100 | Sony | Mouse |
| CD3 | APC/Cy7 | OKT3 | 1:100 | BioLegend | Mouse |
| CD326 (EpCam) | Biotin | 1B7 | 1:200 | eBioscience | Mouse |
| CD4 | BV510 | SK3 | 1:100 | BioLegend | Mouse |
| CD45 | BV605 | 30-F11 | 1:300 | BioLegend | Mouse |
| CD45 | PE | 2D1 | 1:100 | BioLegend | Mouse |
| CD69 | BV711 | FN50 | 1:100 | BD Biosciences | Mouse |
| CD8 | BV421 | RPA-T8 | 1:50 | BioLegend | Mouse |
| CD90 | - | Polyclonal | 1:100 | R&D | Sheep |
| E-Cadherin | - | 24E10 | 1:200 | Cell Signaling Technology | Rabbit |
| GAPDH | HRP | D16H11 | 1:500 | Cell Signaling Technology | Rabbit |
| IL-10Rβ | - | 90220 | 1:200 | R&D | Mouse |
| IL-20Rα | - | 173714 | 1:200 | R&D | Mouse |
| IL-26 | PE | 510414 | 1:50 | Invitrogen | Mouse |
| phospho (Tyr705) STAT3 | - | D3A7 | 1:100 | Cell Signaling Technology | Rabbit |
| Secondary anti-Mouse | Alexa Fluor 488 | Polyclonal | 1:500 | Invitrogen | Donkey |
| Secondary anti-Mouse | Alexa Fluor 647 | Polyclonal | 1:500 | Invitrogen | Donkey |
| Secondary anti-Mouse | HRP | - | 1:500 | Cell Signaling Technology | Horse |
| Secondary anti-Mouse | Cy3 | Polyclonal | 1:500 | Jackson ImmunoResearch | Donkey |

| | | | | | |
|---|---|---|---|---|---|
| Secondary anti-Mouse | Alexa Fluor 555 | Polyclonal | 1:500 | Invitrogen | Donkey |
| Secondary anti-Rabbit | Alexa Fluor 555 | Polyclonal | 1:500 | Invitrogen | Donkey |
| Secondary anti-Rabbit | Alexa Fluor 488 | Polyclonal | 1:500 | Invitrogen | Donkey |
| Secondary anti-Rabbit | HRP | - | 1:500 | Cell Signaling Technology | Goat |
| Secondary anti-Sheep | Alexa Fluor 546 | Polyclonal | 1:500 | Invitrogen | Donkey |
| STAT3 | - | 124H6 | 1:100 | Cell Signaling Technology | Mouse |
| Streptavidin | Alexa Fluor 488 | - | 1:100 | BioLegend | - |
| Streptavidin | PerCP | - | 1:100 | BioLegend | - |
| Streptavidin | BV510 | - | 1:200 | BioLegend | - |
| TCR Vα7.2 | PE/Cy7 | 3C10 | 1:50 | Sony | Mouse |
| TCR Vα7.2 | - | 3C10 | 1:50 | BioLegend | Mouse |
| ZO-1 | - | ZO1-1A12 | 1:100 | Invitrogen | Mouse |

**Table S3 Table of used TaqMan probes**

| **Gene ID** | **TaqMan Probe ID** |
|---|---|
| IL26 | Hs00218189_m1 |
| IL10RB | Hs00175123_m1 |
| IL20RA | Hs01011609_m1 |
| CXCL8 | Hs00174103_m1 |
| IL1B | Hs01555410_m1 |
| TNF | Hs00174128_m1 |
| IFNB1 | Hs01077958_s1 |
